# Perineuronal nets in the rat medial prefrontal cortex alter hippocampal-prefrontal oscillations and reshape cocaine self-administration memories

**DOI:** 10.1101/2024.02.05.577568

**Authors:** Jereme C. Wingert, Jonathan D. Ramos, Sebastian X. Reynolds, Angela E. Gonzalez, R. Mae Rose, Deborah M. Hegarty, Sue A. Aicher, Lydia G. Bailey, Travis E. Brown, Atheir I. Abbas, Barbara A. Sorg

## Abstract

The medial prefrontal cortex (mPFC) is a major contributor to relapse to cocaine in humans and to reinstatement behavior in rodent models of cocaine use disorder. Output from the mPFC is modulated by parvalbumin (PV)-containing fast-spiking interneurons, the majority of which are surrounded by perineuronal nets (PNNs). Here we tested whether chondroitinase ABC (ABC)- mediated removal of PNNs prevented the acquisition or reconsolidation of a cocaine self-administration memory. ABC injections into the dorsal mPFC prior to training attenuated the acquisition of cocaine self-administration. Also, ABC given 3 days prior to but not 1 hr after memory reactivation blocked cue-induced reinstatement. However, reduced reinstatement was present only in rats given a *novel* reactivation contingency, suggesting that PNNs are required for the updating of a familiar memory. In naive rats, ABC injections into mPFC did not alter excitatory or inhibitory puncta on PV cells but reduced PV intensity. Whole-cell recordings revealed a greater inter-spike interval 1 hr after ABC, but not 3 days later. *In vivo* recordings from the mPFC and dorsal hippocampus (dHIP) during novel memory reactivation revealed that ABC in the mPFC prevented reward-associated increases in beta and gamma activity as well as phase-amplitude coupling between the dHIP and mPFC. Together, our findings show that PNN removal attenuates the acquisition of cocaine self-administration memories and disrupts reconsolidation of the original memory when combined with a novel reactivation session. Further, reduced dHIP/mPFC coupling after PNN removal may serve as a key biomarker for how to disrupt reconsolidation of cocaine memories and reduce relapse.

## Introduction

The medial prefrontal cortex (mPFC) is a key region contributing to relapse to cocaine. Output from the medial prefrontal cortex (mPFC) is critical for cocaine-seeking in humans ^1–6^ and reinstatement behavior in rodents ^7–12^. While most studies have focused on mPFC pyramidal neurons, their output is strongly regulated by parvalbumin (PV)-containing fast-spiking inhibitory interneurons ^13–15^.

The majority of mPFC PV neurons are enwrapped in extracellular matrix (ECM) structures called perineuronal nets (PNNs), which facilitate the precise and rapid firing of PV neurons (for review, see ^16^). PV neurons provide dense connectivity to pyramidal neurons in the PFC ^17^ and rapid feedforward inhibition ^15^. PNNs form in the central nervous system during the critical period of plasticity and stabilize neural networks during adulthood ^18^. PNN removal promotes plasticity in adulthood in several memory tasks (for review, see ^19, 20^). However, PNN removal appears to prevent the plasticity induced by strong stimuli, such as drugs of abuse and footshock stress (for review, see ^20, 21^).

We previously demonstrated that degradation of PNNs with the enzyme chondroitinase ABC (ABC) in the prelimbic mPFC, but not the infralimbic mPFC, reduced the acquisition and reconsolidation of cocaine-induced conditioned place preference (CPP) in rats ^22^. Reconsolidation is a memory maintenance process in which memories can be rendered labile upon retrieval (reactivation) and subsequently updated and re-stabilized. Reactivated memories can also be diminished if certain amnestic agents are present during the reconsolidation period (approximately 6 hr after reactivation), including drug-associated memories that are difficult to disrupt ^23–26^. Several studies have shown that, to disrupt stronger memories, *novel information* can be incorporated into the preexisting memory to induce destabilization by the amnestic agent to reduce expression of the old memory ^27–30^. Drug self-administration memories become well established by repeated operant responding for the drug. We therefore tested whether PNN removal in the mPFC could disrupt the reconsolidation of a cocaine self-administration memory by providing novel information at the time of memory reactivation.

PNNs are thought to contribute to the maintenance of memories through their network stabilizing properties ^19, 31^ and may do so through altering firing properties of their underlying PV neurons during the reconsolidation process. PV neurons regulate theta (4-12 Hz) ^38^ and gamma oscillations (30-120Hz) ^28^ and long-range theta/gamma coupling ^36^ . PV neurons support attention ^29, 30^, behavioral flexibility ^31, 32^, and encode reward outcomes ^33^ and expected outcomes ^34^. PV-dependent oscillations are critical to memory ^36, 37^ ^39–41^, and gamma oscillations are altered after cocaine exposure ^32–34^ and when PNNs are removed ^35–41^. Thus, we may expect that removing PNNs would impact the reconsolidation of cocaine memories and also local and long-range oscillations between brain regions. However, only a few studies have measured oscillations across brain regions in addiction models ^42–45^, and no studies have examined how disrupted reconsolidation of a drug memory alters network communication in response to cocaine or cocaine cues. Here we demonstrate that removal of PNNs in the mPFC blocks cue-induced reinstatement and reduces the reinforcing efficacy of a cocaine self-administration memory, but only when a novel contingency is presented at the time of memory reactivation. We further show that PNN removal in the mPFC reduces the synchrony of oscillations between the mPFC and the dorsal hippocampus (dHIP) and alters mPFC cross-hemispheric synchrony.

## Materials and Methods

### Drugs

Cocaine hydrochloride (generous gift from the National Institute on Drug Abuse) was dissolved in saline to intravenously deliver 0.5 mg/kg/infusion in 0.1 mL over 6 sec during all self-administration sessions with exception of the acquisition study, in which 0.125 mg/kg/infusion cocaine was delivered. For the sucrose self-administration experiment, sucrose pellets (Bioserv #F0025) were dispensed for oral consumption during self-administration. Chondroitinase-ABC (ABC; Sigma-Aldrich, C3667) degrades glycosaminoglycan side chains of chondroitin sulfate proteoglycans of PNNs and the “loose” extracellular matrix (ECM) ^46^. ABC was dissolved in sterile water vehicle (Veh) to a final concentration of 0.09 U/µl ^22^.

### Animals

A total of 150 male Sprague Dawley rats were acquired from Simonsen Laboratories (Gilroy, CA), Envigo (Livermore, CA), or bred within the University of Wyoming vivarium. Rats weighed between 275-300 grams at the beginning of the experiment. Animals were pair-housed up until surgery and were then housed singly in a temperature and humidity-controlled room under a 12 hr reverse light/dark cycle, and all experiments were run during the dark cycle. Animals were given food and water *ad libitum* throughout the study with exception of the time they were in the self-administration chambers. All experiments were approved by the Washington State University, University of Wyoming, and Legacy Research Institute Institutional Care and Use Committees, and were done according to the National Institutes of Health *Guide for the Care and Use of Laboratory Animals*. All efforts were made to reduce the number of animals used in the experiments and to minimize pain and suffering.

### Surgery

Rats were anesthetized with isoflurane followed by either intramuscular injection of zyket (ketamine 87 mg/kg; xylazine 13 mg/kg) or intraperitoneal ketdex (ketamine 75 mg/kg; dexmedetomidine 1 mg/kg. Atipamezole (1mg/kg) was given subcutaneously to reverse anesthesia post-operatively. Rats for cocaine self-administration experiments were implanted with chronic indwelling intravenous (iv) catheters. Catheters were made from silastic tubing (RenaSil Silicone Rubber Tubing, .037” OD, 0.025”ID, Braintree Scientific, Inc.) cut to a length of 9.0 cm with a slight taper on one end with a suture bead made from medical grade silicone adhesive (Kwik-sil, World Precision Instruments, Sarasota, FL) placed 2.8 cm from the tapered side for secure anchoring. The right jugular vein was isolated and a small incision was made. The tapered end of the catheter was inserted into the vein until the suture bead was flush with the vein incision. The catheter was secured to the vein by tying 3-0 silk suture thread around the vein and catheter on both sides of the suture bead; additionally, the thread on both sides was tied together. Incisions were sutured, and the animal was given 6-7 days to recover. After surgery, catheters were flushed daily with 0.1 mL of heparin (10 U/mL) and cefazolin antibiotic (10 mg/mL) in sterile saline to help protect against infection and catheter occlusion.

Cannula consisted of either 26-gauge, 1.4 mm spaced, bilateral cannula with dummy obturators (Plastics One, Roanoke, VA) or were made by cutting 26-gauge stainless steel hypodermic tubing (Component Supply Company, Sparta, TN) to a length of 12 mm with 33-gauge stainless steel tubing (Component Supply Company) obturators of the same length as the cannula. Cannula were implanted to target the prelimbic mPFC (+3.2 mm anterior/posterior from bregma, 0.7 mm medial/lateral from midline, and −2.5 mm dorsal/ventral from skull).

For *in vivo* electrophysiology, custom cannulated bilateral tetrode microdrives were implanted in the prelimbic PFC targeting the same site as the cannula above. Tetrodes were just caudal to and lying against the microinjection cannula. A small craniotomy and durotomy was performed over the mPFC and 26-gauge stainless steel cannula of the drive were lowered to 2.5 mm, placing the tetrode guide tubes just above the surface of the brain (for drive details see section on electrophysiology). Three tetrodes per mPFC hemisphere were driven down 2 mm during surgery and were slowly lowered to reach their final depth equal near to where the microinjection would be given. Using a Nano-Z (White-Matter, Inc.), tetrodes were electroplated using Neuralynx gold plating solution (Neuralynx) until all wires had impedances ranging from 200-400 kOhms. To simultaneously record field potentials from the dHIP, 50-micron tungsten wire was lowered into each hemisphere of the dorsal hippocampus (CA1 region, (−4.0 mm anterior/posterior from bregma, 2.2 mm medial/lateral from midline, and −3.5 mm dorsal/ventral from skull). All wires were pinned in place into a 32-channel electrode interface board (Neuralynx). The drives were designed to lower 3 tetrodes per hemisphere to the mPFC and allow for drug delivery as well as record from the mPFC. Qwik-Sil silicone adhesive was used to fill the craniotomy and protect the brain from the dental acrylic used to secure the drive. A stainless-steel ground screw was placed above the somatosensory cortex and a reference screw was placed above the cerebellum.

### Microinjections

Custom-made 33-gauge stainless steel microinjection needles were made to project 1 mm below the cannula and were used to deliver 0.4-0.6 µL of either Veh or ABC over a period of 108 seconds as previously described (Slaker et al, 2015) with a 1 min post-injection period to allow for diffusion. Some groups of animals were naive (***Figures 3, 4, Supplemental Figure 2***) were microinjected during stereotaxic surgery, and guide cannula were not fixed in place as for animals that were used for other experiments. However, the same custom needles, injection volume, and infusion rate were used as with the other animals.

### Self-administration protocol

#### Self-administration training

Animals were trained on one of two different training protocols, including a fixed-ratio 1 (FR1) or fixed-ratio 3 (FR3) schedule of reinforcement. All animals were initially trained during 2 hr sessions on an FR1 schedule in which they were placed in operant chambers (Med Associates, Fairfax, VT) with one active (right lever) and one inactive (left) lever, with a cue light over each lever and a house light on the wall opposite from the levers. Active lever presses were reinforced with a 0.1 mL infusion of cocaine (either 0.125 mg/kg/infusion for the acquisition experiment or 0.5 mg/kg/infusion for all other experiments) or a 45 mg sucrose pellet (Bio-Serv, Flemington, NJ). Reward delivery was accompanied by simultaneous cue light activation for 6 sec and house light activation for 20 sec that signaled a time-out period during which active lever pressing produced no consequences. Inactive lever presses at any time had no consequences. All active and inactive presses were recorded. To be included in the study, animals had to reach three consecutive days with >10 rewards. For FR3 training in a subset of animals, rats began on an FR1 schedule and had to reach the criterion above before being switched to an FR3 schedule in which the animals had to press the active lever three times to be rewarded a single cocaine infusion, cue light, and given the time-out light. FR3-trained animals, in addition to meeting the FR1 to FR3 transition criteria, had to have three or more days of FR3 training within the total 10 training sessions. 3-4 hr following the last training session, most cohorts received a microinjection of either vehicle (Veh) or ABC targeted to the prelimbic mPFC. One additional cohort received Veh or ABC targeted to the prelimbic mPFC within approximately 1 hr after the memory reactivation session (see below).

#### Memory reactivation and cue reinstatement

Three days after the last training session, FR1 trained animals received either a 30 min FR1 or a novel variable ratio schedule (VR5 reactivation session). VR5 reactivation consisted of animals responding to a VR5 schedule of reinforcement in which the number of active presses needed to receive each reward was randomly selected from 1-9 presses with an average of 5.

One set of FR1- trained animals was not microinjected following training but was instead microinjected *after* the VR5 reactivation session, with the Veh or ABC microinjection occurring within a 5 min to 2 hr window for all but one animal, which received a Veh microinjection 2.4 hr after the VR5 memory reactivation session. The average time (± standard error of the mean, SEM) for post-VR5 microinjection was 67 ± 10 min after the completion of the 30 min memory reactivation session in this group. Hereafter, this group is referred to as “post-VR5” (***Figure 2***). For naive animals receiving microinjections of Veh or ABC but no self-administration training, we refer to the groups as either “1 hr”, since infusions took place approximately 1 hr after memory reactivation, or “3 days”, both of which were the interval between microinjection and experimental endpoints taken (***Figures 3 and 4***; ***Supplemental Figure 2***).

The FR3 trained animals received a 30 min FR3 reactivation session. One set of animals, a no reactivation group, did not receive a reactivation session but received Veh or ABC in their home cages. The day following the reactivation session, *all animals* received a 30 min extinction session in which no cues or rewards were delivered. Immediately following the 30 min extinction, animals underwent a 30 min cue-induced reinstatement session in which a single cue light primed animals to continue to press on an FR1 schedule for the cue light above the cocaine lever, but no rewards were delivered (cocaine or sucrose). The day following the extinction and cue-reinstatement sessions, most groups of rats received three consecutive days of a 3 hr long progressive ratio (PR) self-administration in which the reinforcement schedule needed to obtain a reward increased as previously described ^47^. Following the PR days, rats were euthanized with an overdose of sodium pentobarbitol (67 mg/kg, ip) and intracardially perfused with 4% paraformaldehyde. The brains were removed and cryoprotected in 20% sucrose and were frozen with powdered dry ice and stored at −80 °C until analysis for cannula placement and confirmation of PNN degradation.

### BS3 crosslinking and western blot

For the crosslinking surface protein quantification assay, we followed the procedure of Boudreau ^48^, with slight modifications. In brief, following cocaine self-administration training, one group of rats underwent the FR1 reactivation protocol and the other group underwent the VR5 reactivation protocol described above. Immediately following the 30 min reactivation sessions, each animal was decapitated and the brain was extracted. A ∼2 mm coronal section was taken encompassing either the mPFC or the amygdala. The section was then placed on a piece of wetted filter paper placed on an inverted glass petri dish on ice, and the brain region of interest was rapidly dissected with a razor blade. The dissected region was rapidly sliced into strips of ∼400 microns by hand before being transferred to a sample tube containing 1 mL of crosslinking solution. The crosslinking solution was composed of ice-cold artificial cerebral spinal fluid (aCSF) (1.2 mM CaCl_2_, 20 mM HEPES, 147 mM NaCl, 2.7 mM KCl, 1 mM MgCl_2_, 100 mM dextrose in ddH_2_O) mixed with 52 mM bis(sulfosuccinimidyl)suberate (BS^3^; ThermoFisher Scientific, catalog 21580), a membrane-impermeable bifunctional crosslinking agent, in 5 mM sodium citrate buffer (5 mM citric acid and 5 mM sodium citrate, pH 5.0) to bring the final concentration of BS^3^ to 2 mM in aCSF. Following transfer to the tube of crosslinking agent, the sample tube was incubated at 4°C for 30 min gently mixing on a rocker. The crosslinking reaction was quenched by adding 100 uL of 1 M glycine to the sample tube, which was then placed on the rocker for an additional 10 min at 4°C. After quenching, the samples were centrifuged at 20,000 x g for 2 min at 4°C. The supernatant was discarded and the pellet was resuspended in 400 uL of NP-40 lysis buffer (pH 7.4, 25 mM HEPES, 500 mM NaCl, 2 mM EDTA, 1 mM DTT, 0.1% Nonidet P-40 (v/v) with phosphatase inhibitors (Pierce, A32957 or Millipore-Sigma, 4906845001) and EDTA-free protease inhibitors (Pierce, A329965). Samples were then sonicated for 5 sec before centrifuging for 2 min at 20,000 x g at 4°C. The supernatant was aliquoted into 20 uL aliquots and flash frozen on dry ice and stored at −80°C. A BCA protein assay kit (ThermoFisher, catalog number: 23225) was used to determine protein concentration. 40 µg of sample protein was diluted with NP-40 buffer and SDS Sample Buffer (Bio-Rad) and were then loaded into 10% precast gels (Bio-Rad) and electrophoresed with Tris/Glycine/SDS running buffer (Bio-Rad) before being transferred to PVDF membranes. The blots were then washed 3 times for 5 min in TBS-T (10mM Tris/HCL, pH 8.0, 15 M NaCL, 0.05% Tween 20) before being blocked for 1 hr in Odyssey blocking buffer (LI-COR, Inc.). The blots were then probed with primary antibodies (GluN2B 1:500 Abcam, 71-8600) in Odyssey blocking buffer on a shaker at 4°C overnight. Blots were then washed in TBS-T 3 times for 10 min before incubating at room temperature on a shaker in HRP conjugated secondary antibody solution (1:10,000) in blocking buffer. After washing in TBS-T 3 more times for 10 min each, the blots were incubated in western blotting substrate (ThermoFisher Scientific, 34579) for 2 min and imaged on a ChemiDoc Touch Imaging System (Bio-Rad). To normalize to total protein concentration, blots were incubated in Ponceau S staining solution (Sigma-Aldrich) for 5 min to detect protein before being imaged on the ChemiDoc Touch. Each protein band was normalized to the total protein intensity of the entire lane.

### Immunohistochemistry

#### Analysis of PNNs and PV at 1 hr and 3 days

To assess the intensity and number of PV and PV/WFA cells at 1 hr and 3 days after ABC microinjection, animals were anesthetized with pentobarbital (67 mg/kg, ip) and perfused intracardially with 4% paraformaldehyde (PFA) in phosphate buffered saline (PBS). Brains were removed and stored overnight in 4% PFA at 4 °C. The following day, brains were moved to a 20% sucrose solution, and 24 h later were frozen with powdered dry ice and stored at −80 °C until analysis. Coronal sections (40 µm) through the mPFC were sectioned using a sliding microtome and stained with WFA (1:500, Vector Laboratories, FL-1351, RRID AB_2336875) and rabbit anti-PV antibody (1:1000, Novus, NB120-11427, RRID AB_791498) according to ^49^. For each rat, we analyzed 3-5 images/hemisphere/rat (1 rat had only one image). PNNs were identified by staining with *Wisteria floribunda* agglutinin (WFA), a plant lectin that binds to chondroitin sulfate proteoglycan components of the PNNs ^50^. PNN “stubs” after ABC digestion were identified by staining with mouse anti-chontroitin sulfate (1B5) antibody (1:20, Amsbio, 270431-CS). Secondary antibodies were AlexaFluor 405 goat anti-rabbit for PV or AlexaFluor 594 goat anti-mouse (1:500, ThermoFisher Cat# A31556).

The mPFC and dHIP images were taken using a Leica SP8 laser scanning confocal microscope with an HCX PL Apo CS2, dry, 20X objective with 0.75 numerical aperture (NA) or with a water immersion 63X objective with 1.2 NA. Images (0.84 µm/pixel for 20X, 0.23 µm/pixel for 63X) were compiled into summed images using ImageJ macro plug-in Pipsqueak ^51^ or PipsqueakTM, scaled, and converted into 8-bit, grayscale tiff files. Pipsqueak ^51^ or PipsqueakTM was run in “semi-automatic mode” to select ROIs to identify individual PV cells and PNNs. The plug-in compiles this analysis to identify single, double-labeled, and triple-labeled neurons (https://ai.RewireNeuro.com). Histologies for targeted sites and for PNN loss in ABC-treated rats were identified by at least two investigators blinded to experimental treatment.

#### Analysis of puncta around PV cells at 1 hr and 3 days

Immunohistochemical methods and antibodies used for this study were similar to those described previously ^52^. Confocal imaging was performed on either a Zeiss LSM 780 or a Zeiss LSM 900 confocal microscope with a 63 x 1.4 NA Plan-Apochromat oil objective (Carl Zeiss MicroImaging, Thornwood, NY) due to a change in instrumentation in the OHSU Advanced Light Microscopy Core mid-way through the study. Coronal mPFC tissue sections from a particular animal were only imaged on one of the microscopes. The imaging strategy was similar to what was described previously ^49, 53^. The dimensions of images captured using the LSM 900 confocal microscope (xyz: 126.77 x 126.77 x 0.230 µm) were smaller than those captured on the LSM 780 (xyz: 134.95 x 134.95 x 0.384 µm) due to improved pixel resolution in all planes. The change in Z-axis pixel thickness affected the number of optical slices used to create the Z-stack subsets through the middle of each PV neuron. Using Zen software (Carl Zeiss, RRID SCR_013672), PV neurons that met our criteria for inclusion in the study were identified.

Inclusion criteria for PV neurons included being fully within the boundaries of the field of view and having a visible nucleus within the Z-stack subset. For LSM 780 confocal images, three consecutive 0.384 µm optical slices were used for each Z-stack subset for a total thickness of 1.152 µm through the middle of each PV neuron. For LSM 900 confocal images, five 0.230 µm optical slices were used for each Z-stack subset for a total thickness of 1.15 µm. Image analysis of GAD65/67- and VGluT1-labeled puncta apposing the exterior boundaries of the PV neuron was performed using Imaris image analysis software as previously described ^49, 52^. All parameters to identify puncta with the Imaris Spots segmentation tool were the same, regardless of the confocal microscope used to image that particular tissue section.

### Whole-cell patch clamp electrophysiology

Whole-cell patch clamp recordings of fast-spiking interneurons (FSIs) were performed, as FSIs are highly likely to contain PV ^54^. Recordings in brain slices from adult (P60-P90) male rats with and without PNNs in layer V of the prelimbic mPFC were done using established procedures ^49, 53^. Rats were microinjected with either Veh (0.6 µL saline) or ABC (0.09 U/µL, 0.6 µL/hemisphere) bilaterally into the prelimbic mPFC (AP +3.2, DV −2.5, ML ± 0.8) over 2 min and were given 1 min to be absorbed before removing the cannula. 1 hr or 3 days after microinjection, rats were perfused, decapitated, and coronal slices (300 µm) containing the mPFC (2.8-3.5 mm from bregma) were prepared using a vibratome (Leica VT1200S). Brains were perfused and cut in a choline-based cutting solution (in mM): 2.5 KCl, 84.01 NaHCO_3_, 119

NaH_2_PO_4_, 180 D-glucose, 0.5 CaCl_2_, 7 MgCl_2_, 110 choline Cl, 3 Na-pyruvate, 11.6 Na- ascorbate, 300-310 mOsm, pH 7.2-7.3. After cutting, slices were incubated for 10 min in artificial cerebrospinal fluid (aCSF) at 32°C followed by a minimum of 1 hr in aCSF prior to recording (in mM): 120 NaCl, 2.5 KCl, 26.2 NaHCO_3_, 1.25 NaH_2_PO_4_, 11 Glucose, 2.5 CaCl_2_, 1.3 MgCl_2_, 300-310 mOsm, pH 7.2-7.3.

Prior to recording, slices were incubated with WFA (1 µg/mL) in aCSF for 5 minutes to stain for or verify reduction of PNNs. Fluorescing neurons positive for WFA in the vehicle group were identified in layer V of the prelimbic mPFC using CellSens software (Olympus) and subsequently patched. For rats microinjected with ABC, WFA stain was used to verify a reduction in positive WFA neurons and FSIs were visually and electrophysiologically identified. For intrinsic experiments, 1.5-mm borosilicate glass (WPI, Sarasota, FL) pipettes were filled with an intracellular solution containing (in mM) 145 K gluconate, 3 KCl, 2 MgCl_2_, 2 Na^+^ - ATP, 10 HEPES, 0.2 EGTA, pH 7.2-7.3 adjusted using potassium hydroxide, and 290-300 mOsm. FSIs in layer V of the prelimbic mPFC were identified as described above and with their response to current injection (−100 to 500 pA, 100 pA increments, 250-500 ms) using previously established criteria ^53^. Elicited action potentials were recorded, counted, and analyzed using pClamp11.0.2 (Clampfit, Axon Instruments, Sunnyvale, CA). Neuronal excitability was assessed by quantifying the number of action potentials generated in response to each depolarizing current injection.

Intrinsic properties were collected from the first action potential elicited at 200 pA current injection. Input resistance was determined by the change in voltage from −100 pA current injection. Rheobase is the minimum amount of current to elicit an action potential. The action potential threshold was determined using the maximum second derivative method^55^. The peak amplitude (difference between the AP peak and threshold) and amplitude of the afterhyperpolarization (AHP) were measured from the calculated action potential threshold.

Capacitance was measured from the membrane properties within pClamp. The interspike intervals (ISIs) were calculated using Clampfit 11.2 using the threshold search function, with the baseline set at −70 mV and the threshold for spike detection set at 0 mV. The values for ISIs from the 200 pA and 500 pA traces were taken, and the mean and standard deviation were calculated. The coefficient of variation (CV) was then calculated by dividing the standard deviation by the mean for each trace from each animal.

### *In vivo* electrophysiology

#### Recording and LFP analysis

For recording, 32-channel headstages (Intan Technologies) were connected to a motorized commutator (Plexon) and an OpenEphys data acquisition system. All channels were referenced to a screw electrode placed over the cerebellum (reference shorted to ground), filtered between 1 and 7500 Hz, multiplexed, and digitized at 30 kHz. A Med-Associates TTL-output card was used to deliver TTL pulses to the digital inputs of the OpenEphys data acquisition system to timestamp lever presses, cue-lights, and houselights used in the self-administration paradigm.

#### Local field potential (LFP) analyses

All electrophysiology analyses were performed using custom Python scripts. Signals were referenced to a screw electrode over the cerebellum, downsampled from 30 to 2 kHz, and bandpass filtered between 1 and 300 Hz with a Butterworth bandpass filter using *butter* and *filtfilt* functions from the *signal* module of Python’s scientific library *scipy.* Lever press timestamps were recorded via TTL pulses and 2.5 s sections of data centered around each TTL pulse were sliced and considered as trials. A 2.7 root mean square (rms) threshold was set for artifact detection, and any section of data which exceeded this threshold was considered artifactual. Trials containing any artifacts were automatically rejected. Following artifact detection, trials were manually curated by visual inspection of each trial; trials with sections that were clearly artifactual but remained undetected by rms thresholding were removed.

Spectrograms were calculated via Morlet wavelet convolution from 1 to 120 Hz, implemented using the *fft* and *ifft* functions included in python’s numerical processing library, *numpy*. The number of wavelet cycles varied from 3 to 9, increasing as the frequency of the wavelets increased from 1 to 120 Hz, linearly. Power was extracted by taking the square of the real component of the analytic signal. A baseline period was defined from −1000 ms to −100 ms, where each press occurred at 0 ms. Each trial was normalized at every frequency by dividing each sample by the mean power of the baseline period. Spectrograms were then generated by taking the average of all trials. Spectrograms were quantified at various frequency bands by defining a time-frequency window; then for each trial, power was averaged across the time-frequency window and statistical significance was assessed via two-way ANOVAs.

Phase-amplitude coupling was assessed by calculating both mean vector length (MVL) as described ^56^, and modulation index (MI) as described ^57^. To generate comodulograms, phase angles from 3 to 13 Hz and amplitude from 20 to 100 Hz were extracted via Morlet wavelet convolution for each trial as described above. MVL was calculated from the concatenated trials for every pair of phase-amplitude frequencies. Each MVL was then z-scored by permutation testing with 10,000 permutations, where each permutation was generated by time-shifting the amplitude timeseries of each trial by a randomly generated number of samples before concatenation. The MVL was calculated for each permutation and the observed MVL was standardized to the distribution of the permuted MVLs with a significance threshold set at z = 1.96. To confirm the coupling between frequencies indicated by the z-scored comodulograms, gamma power and theta phase timeseries were extracted via a filter-Hilbert method using a Butterworth bandpass filter and the *Hilbert* function from Python’s *scipy* library. Trials were concatenated and phase was binned into 30 equally sized bins. The mean normalized power in each phase bin was taken to generate a distribution of power over phase. The MI is calculated from this distribution, and then z-scored by permutation testing with 10,000 permutations, where each permutation was generated by randomly shuffling trial IDs before concatenation. The MI was calculated for each permutation and the observed MI was standardized to the distribution of the permuted MIs with a significance threshold set at z = 1.65.

#### Lag analysis and directionality

The directionality of signals across different sites can be assessed directly by calculating the cross-correlation of amplitude envelopes between the two sites at varying lags in a sliding window fashion, as described ^58^. First, envelopes of instantaneous theta amplitude (4-12 Hz) from the signals of two different sites were computed by bandpass filtering signals from 4 to 12 Hz, applying the Hilbert transform, and then taking the magnitude of the real component of the resulting analytic signal. The envelope from one site was incrementally shifted from −100 ms to +100 ms. At each increment, Spearman’s rho was calculated between the envelopes of the two sites. Correlation coefficients were normalized to the minimum and maximum observed correlation within each trial and the mean normalized correlation +/- standard error of the mean (SEM) was plotted for each lag. For each trial, the lag at which peak cross-correlation occurs was taken to generate a distribution of peak lags. Statistical significance between Veh and ABC treatment was assessed with the Kolmogorov-Smirnov test. To determine whether a significant lead was observed, for each trial, normalized correlation coefficients were averaged across all negative lags and all positive lags (zero excluded) and statistical significance was assessed with a two-way ANOVA. If the mean normalized correlation coefficient of all negative lags is significantly different from the mean normalized correlation coefficient of all positive lags, then it must be the case that signals from one region lead the other, and directionality is denoted by the sign of the lag. Note that in the current work, a negative lag denotes that the dHPC leads the mPFC; additionally, in the supplemental figures, a negative lag denotes that the left hemisphere of the mPFC leads the right hemisphere of the mPFC.

Like lag cross-correlation analysis, Granger Causality (GC) testing can also be used to assess the directionality of the flow of information between to areas. Given two time series, 𝑋(𝑡) and 𝑌(𝑡), if information about the past of 𝑋(𝑡) and 𝑌(𝑡) is more useful in making predictions about future values of 𝑋(𝑡) than information about the past of 𝑋(𝑡) alone, we may say that 𝑌(𝑡) Granger causes 𝑋(𝑡) in the 𝑌 → 𝑋 direction. Mathematically, we compared two models representing these data: a univariate autoregressive (AR) model built only on data from 𝑋(𝑡), and a bivariate AR model built on data from both 𝑋(𝑡) and 𝑌(𝑡). If the error term from the bivariate AR model 𝑒_()_, is smaller than the error term of the univariate AR model 𝑒_(_; that is, information from both time series is a better predictor of 𝑋(𝑡) than just information from 𝑋(𝑡) alone, then we may consider that 𝑌(𝑡) Granger causes 𝑋(𝑡), and so GC is given by 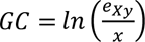^59^.

To ensure stationarity, signals were first detrended and normalized by subtracting ERPs and z-scoring. Next, univariate and bivariate autoregressive models were generated by a custom Python implementation of MATLAB’s *armorf* function. Models were built on data from a 400 ms time window, with a model order of 6. Optimal model order was selected by minimizing Bayesian information criterion. This process was done in a sliding window fashion at increments of 0.01 s to calculate GC across time for each trial. Time domain GC was normalized to a baseline period and statistical significance was assessed via two-way ANOVA. For a one-tailed test, p ≤ 0.05 if Z ≥ 1.65; for two-tailed test, p < 0.05 if Z ≥ 1.96.

### Statistical analyses

All statistical data with exception of some of the *in vivo* electrophysiology studies (see above) were analyzed using GraphPad Prism 9.5.1. Behavioral data were analyzed with a two-way analysis of variance (ANOVA), a two-way ANOVA with repeated measures over time or day (RM-ANOVA), or a three-way repeated measures ANOVA with a Greenhouse Geisser correction followed by Šídák’s post-hoc test, with significance set at p < 0.05. In the case of two comparisons between treatment groups, a Student’s unpaired, two-tailed t test was conducted. For analysis of WFA or PV intensity in the immunohistochemical studies, and for analysis of relative power in the spectrograms (***Figure 5***), due to non-normal distribution, data were log transformed and then subject to analysis using a two-way ANOVA. Puncta data were analyzed using a two-way ANOVA followed by a Šídák’s post-hoc test. For analysis of neuronal excitability, data were analyzed with a three-way, RM ANOVA. For analysis of firing variability and intrinsic properties, data were analyzed with a two-way, RM ANOVA followed by a Šídák’s post-hoc test to compare significant interactions. All results are summarized as mean ± SEM unless otherwise stated. Differences were considered significant if p < 0.05 and trends are reported if p < 0.06. Statistics used for *in vivo* electrophysiology are described separately within each paragraph under “EEG Recordings” above. All statistics are reported in ***Supplemental Tables 4-11***, and statistically significant differences are reported in the figure legends.

## Results

### ABC in the mPFC blunts acquisition of cocaine self-administration

We tested whether ABC in the mPFC would suppress the acquisition of cocaine self-administration using a low dose of cocaine previously used for acquisition studies ^60^. ***Figure 1A*** shows the experimental timeline, and the sites targeted for Veh or ABC microinjection are shown in ***Figure 1E*.** Rats increased the number of active lever presses and cocaine infusions across the 20 days of training. However, ABC treatment in the mPFC reduced the number of active presses and infusions compared with the Veh group (***Figures 1B, 1C; Supplemental Table 1 for total lever presses and rewards; Supplemental Table 2 for statistics***). There were no effects of PNN removal for inactive lever presses (***Figure 1D***; ***Supplemental Table 2***). Thus, the acquisition of a cocaine-associated memory was blunted after PNN degradation in the mPFC.

**Figure 1.**
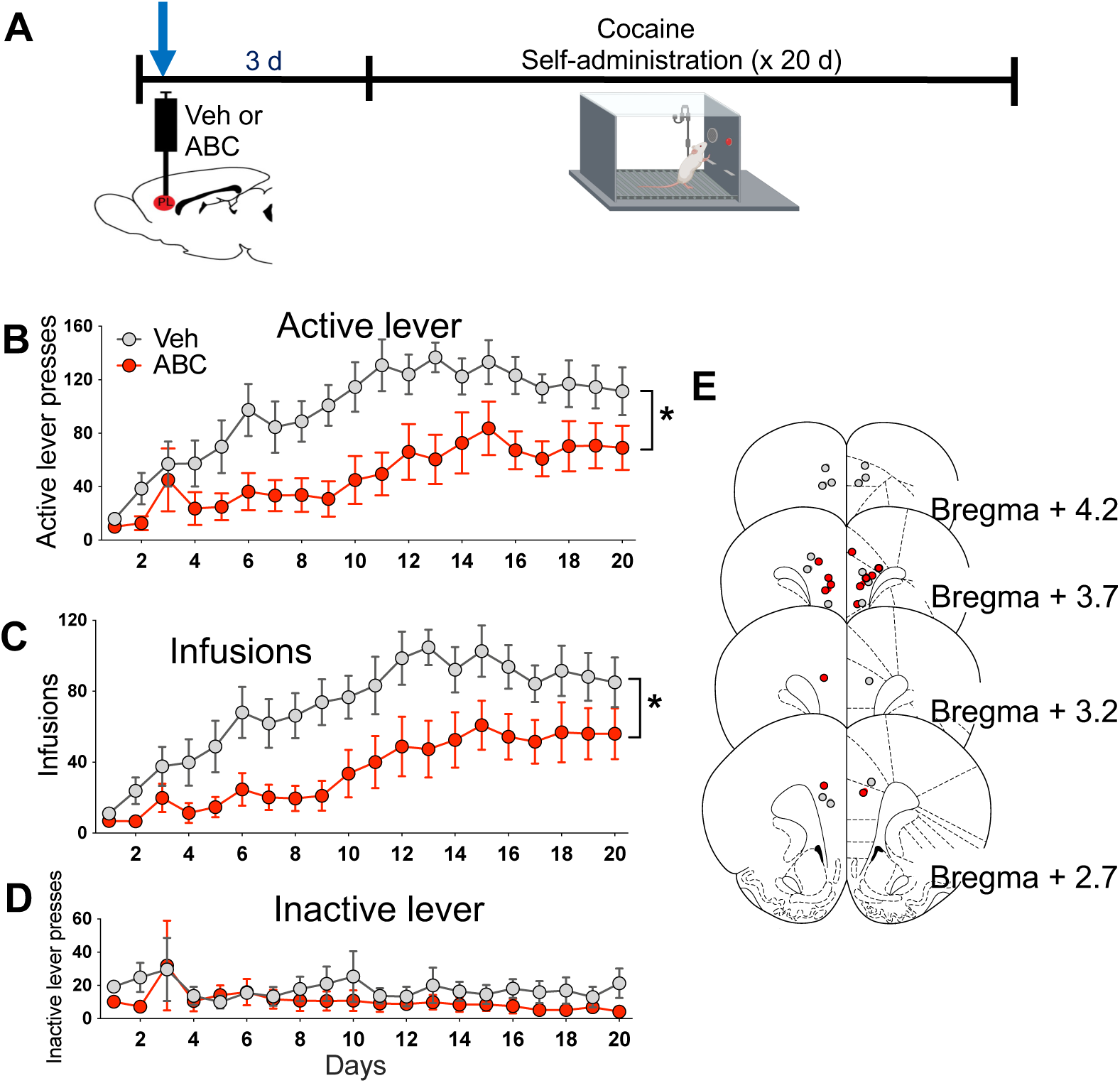
ABC treatment in the mPFC blunts acquisition of cocaine self-administration. **(A)** Timeline of experiment. **(B)** Active lever presses during acquisition are reduced after ABC treatment compared with vehicle (Veh) treatment. A two-way repeated measures (RM) ANOVA showed a main effect of treatment (F_1,12_ = 11.10, p = 0.006), and a main effect of day (F_3.22,38.58_ = 8.815, p < 0.0001). **(C)** Number of cocaine infusions during acquisition are reduced after ABC treatment. Two-way RM ANOVA showed a main effect of treatment (F_1,12_ = 9.167, p = 0.0105) and a main effect of day (F_2.90,31.14_ = 10.71, p < 0.0001). **(D)** Inactive lever presses during acquisition were not different between groups. **(E)** Targeted microinjection sites in the dorsal PFC. Gray dots = Veh; red dots = ABC. Data are mean ± SEM. Veh N = 8; ABC N = 6. + p < 0.05 compared with Day 1 within groups; * p < 0.05 compared between groups on the same day.

### ABC in the mPFC blocks cocaine cue-induced reinstatement when given prior to a novel VR5 reactivation session

We next tested whether ABC treatment in the mPFC *after* training but prior to or after memory reactivation would alter reactivation or subsequent reinstatement. Because it was previously found that cocaine self-administration memories may not be updated unless a novel contingency is introduced during memory reactivation ^27^, we also examined the effect of ABC treatment when the reactivated memory was updated with a *novel VR5 schedule*. ***Figure 2A*** shows the experimental timeline, and ***Figures 2L, M*** show the sites for Veh and ABC microinjection, which spanned prelimbic and dorsal anterior cingulate regions. ***Figures 2B and 2C*** show there were no late training differences among groups in active lever presses (***Figure 2B***), number of cocaine infusions (***Figure 2C***), or inactive lever presses (***Supplemental Tables 3 and 4*).** Rats received Veh or ABC microinjection into the mPFC either 3 days prior to or within approximately 1 hr after the memory reactivation session (Pre-VR5 or Post-VR5, respectively). During the memory reactivation session, there was no effect of ABC treatment on active lever presses, infusions, or inactive lever presses, irrespective of FR1 or VR5 schedules. ***(Figures 2D, 2E; Supplemental Tables 3 and 4***).

**Figure 2.**
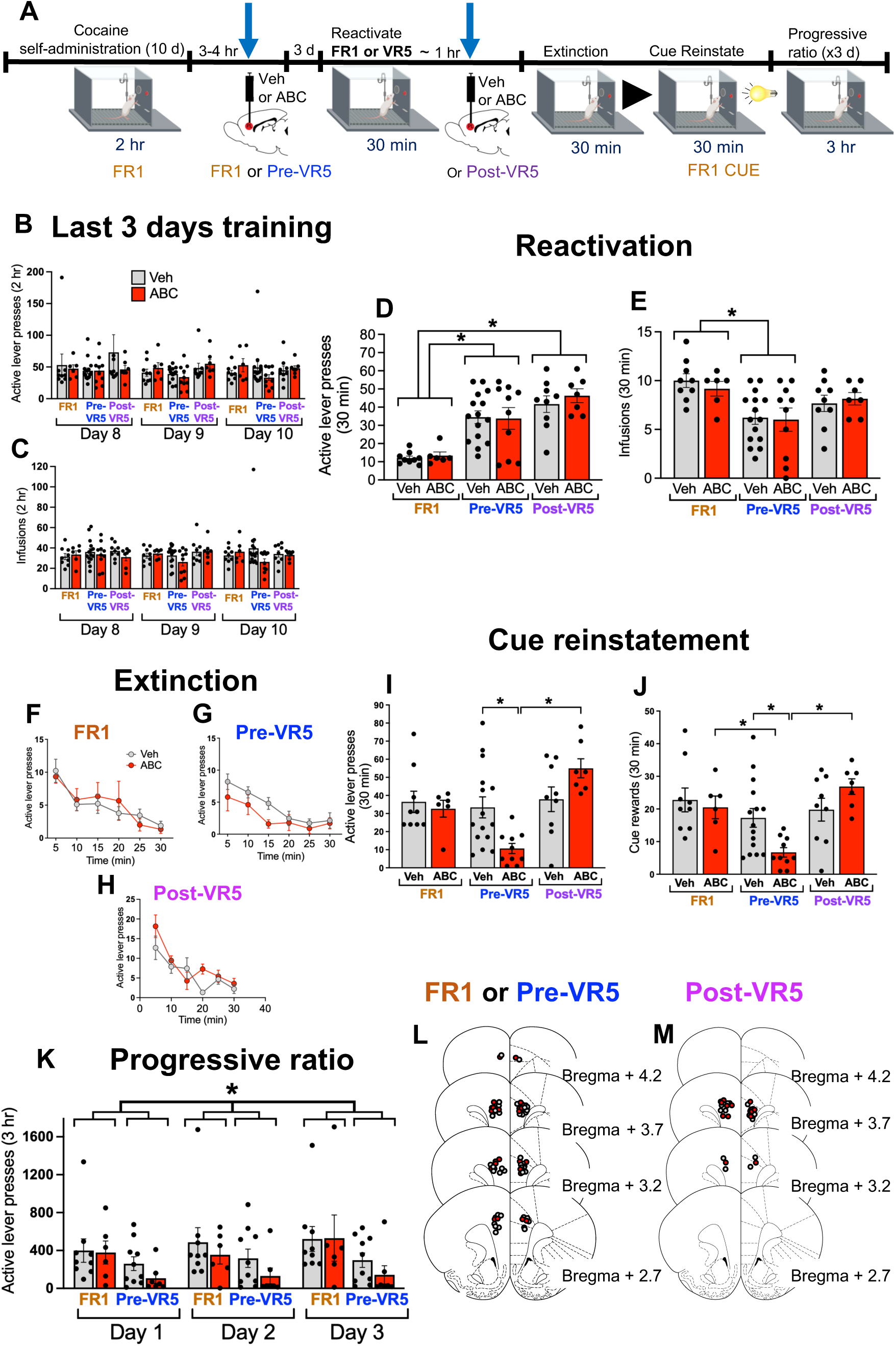
ABC treatment in the mPFC blocks cocaine cue-induced reinstatement when given prior to a novel VR5 memory reactivation session. **(A)** Timeline of experiment. **(B)** Active lever presses during the last 3 days of training. All rats were trained on an FR1 schedule. A three-way RM ANOVA showed no differences among groups. **(C)** Number of cocaine infusions during the last 3 days of training. A three-way RM ANOVA showed no differences among groups. **(D)** Active lever presses during memory reactivation were increased in the VR5 compared with FR1 groups. A two-way ANOVA showed a main effect of reactivation type (F_2,50_ = 23.98, p < 0.0001) but no main effect of ABC treatment. In a subsequent one-way ANOVA and Šídák’s post-hoc test, active lever presses were lower in FR1 vs. Pre- and Post-VR5 groups (p< 0.0001 for both). **(E)** Number of cocaine infusions during memory reactivation. A two-way ANOVA showed a main effect of reactivation type (F_2,50_ = 8.136, p = 0.009), and a one-way ANOVA followed by a Šídák’s post-hoc test showed that infusions were higher in the FR1 vs Pre-VR5 group (p = 0.0003). **(F-H)** Extinction of active lever presses in FR1, Pre-VR5, and Post- VR5 groups, respectively. There were no differences between treatment groups, with only an effect of time in all three groups (FR1 reactivation: F_3.196, 41.54_ = 7.467, p = 0.0003; Pre-VR5 reactivation: F_3.583, 82.40_ = 9.779 p < 0.0001; Post-VR5 F_3.081, 43.13_ = 13.79, p < 0.0001). **(I)** Active lever presses during cue reinstatement. A two-way ANOVA showed a main effect of reactivation type (F_2,50_ = 9.303, p = 0.0004) and a treatment x reactivation interaction (F_2,50_ = 6.145, p = 0.0041). Pre-VR5 ABC-treated rats had decreased lever pressing vs. Veh controls (p = 0.0079) and vs. Post-VR5 ABC rats (p < 0.0001). **(J)** Number of cue rewards during cue reinstatement, showing a main effect of reactivation type (F_2,50_ = 8.612, p = 0.0006) and a treatment x reactivation interaction (F_2,50_ = 4.266, p = 0.0195). Pre-VR5 ABC-treated rats had a decreased number of cue rewards vs. Veh controls (p = 0.0233) vs. FR1 ABC-treated rats (p = 0.0167), and vs. Post-VR5 ABC-treated rats (p = 0.0002). **(K)** Active lever presses during progressive ratio (PR) schedule of reinforcement over three consecutive days in FR1 and Pre-VR5 groups. A three-way RM ANOVA showed a main effect of reactivation (F_1,28_ = 4.665, p = 0.039) across days, with a higher reactivation in the FR1 vs Pre-VR5 group (Post-VR5 group was not tested). (**L, M)** Microinjection sites for FR1 or Pre-VR5 **(L)** or Post-VR5 **(M)**. Gray dots = Veh, red dots = ABC. Data are mean ± SEM. FR1: Veh N = 9; ABC = 6; Pre-VR5: Veh N = 15 (10 for PR); ABC N = 10 (7 for PR); Post-VR5: Veh N = 9; ABC N = 7. * p < 0.05.

During extinction, there were no differences in active lever presses between Veh- and ABC-treated rats in any group, with decreased responding over session time (***Figures 2F-H***). During cue reinstatement, ABC treatment prior to VR5 (but not FR1) memory reactivation reduced active lever presses and associated rewards during a 30 min cue reinstatement session. ABC treatment *after* memory reactivation had no effect, irrespective of FR1 or VR5 schedule (***Figures 2I, 2J***).

Rats were subsequently tested on a Progressive Ratio (PR) schedule of reinforcement to determine whether 1) reduced responding in Pre-VR5 ABC-treated animals would persist for additional days, and 2) the reinforcing efficacy of cocaine was altered ^47^ when rats could once again obtain cocaine infusions. There was a main effect of reactivation type, with reduced active lever pressing in the Pre-VR5 vs. FR1 group (***Figure 2K***) and a similar reduction in cocaine infusions and inactive lever presses in the Pre-VR5 vs. FR1 group (***Supplemental Tables 3 and 4***). Thus, PNN removal reduced cue responding only when a novel VR5 reactivation session was combined with removal of PNNs in the mPFC prior to VR5 memory reactivation, suggesting that PNNs are necessary for updating or maintaining stability of the cocaine memory.

### ABC in the mPFC does not block cocaine cue-induced reinstatement in the absence of memory reactivation, nor does it block sucrose cue-induced reinstatement

We examined three additional control groups given Veh or ABC treatment. The first group was a “no reactivation” group that did not receive memory reactivation. The second group self-administered cocaine on an FR3 schedule of reinforcement to test whether higher active lever pressing during the VR5 memory reactivation session (vs. the FR1 group; see ***Figure 2D***) could account for better memory recall during reactivation and therefore reduce subsequent cue reinstatement in the ABC treatment group. The third group self-administered sucrose rather than cocaine to determine whether ABC combined with VR5 reactivation would alter responding for a non-drug reward. We used the same timeline as in ***Figure 2A***, with Veh or ABC given 3 days prior to the reactivation session. Brain sites for microinjection are shown in ***Supplemental Figures 1R, 1S***. There were no late training differences among groups in active lever presses (***Supplemental Figures 1A-D)***, nor did treatment with ABC alter reactivation, extinction, cue-induced reinstatement, or responding on the progressive ratio schedule (***Figures 1E-Q; Supplemental Tables 5 and 6***).

### ABC treatment reduces PV cell immunolabeling intensity

Given the dependence of PNN removal prior to memory reactivation but not after reactivation on subsequent reduction in cue-reinstatement, experiments were done to identify whether PNN removal at different timepoints would lead to changes in PV neurons. We determined in naive rats whether there were changes in PV staining intensity levels when ABC was delivered 3 days prior vs. approximately 1 hr after memory reactivation, the latter of which occurred in the post- VR5 reactivation group. ***Figures 3A*** and ***3B*** show WFA-labeled PNNs in naive rats given unilateral microinjections of Veh or ABC into the mPFC. ABC decreased WFA staining intensity levels of not only PNNs but also apparent reduction of WFA-stained loose ECM. This more dorsal prelimbic target caused some spread into the dorsal anterior cingulate. ***Figure 3C*** shows a double-labeled PV/WFA cell, and ***Figure 3D*** shows a double-labeled PV/Stub antibody cell. Stub antibody detects remnants of PNNs digested with ABC ^61^. Since not all PV cells have PNNs, we assessed the intensity and number of total PV cells, PV cells that were surrounded by WFA after Veh treatment, and PV cells that were surrounded by Stub antibody. ABC treatment decreased PV intensity at both time points when all PV cells were assessed (***Figure 3E***), while the total *number* of PV cells was not altered (***Figure 3F***). The intensity of PV in double-labeled PV neurons surrounded by either WFA (Veh side) or Stub antibody (ABC side) was reduced at 3 days, but not 1 hr, after ABC treatment (***Figure 3G***), together indicating that 3 days after ABC treatment, PV intensity was more affected in cells that were not originally surrounded by PNNs. The *number* of PV cells surrounded by PNNs showed a strong trend toward a decrease at both timepoints (p = 0.0625) (***Figure 3H***). Altogether, these results show that ABC decreased the intensity of PV in neurons surrounded by PNNs 3 days later, but not 1 hr later, consistent with others demonstrating a decrease in PV intensity after PNN removal ^62–64^ (statistical data shown in ***Supplemental Table 7***).

**Figure 3.**
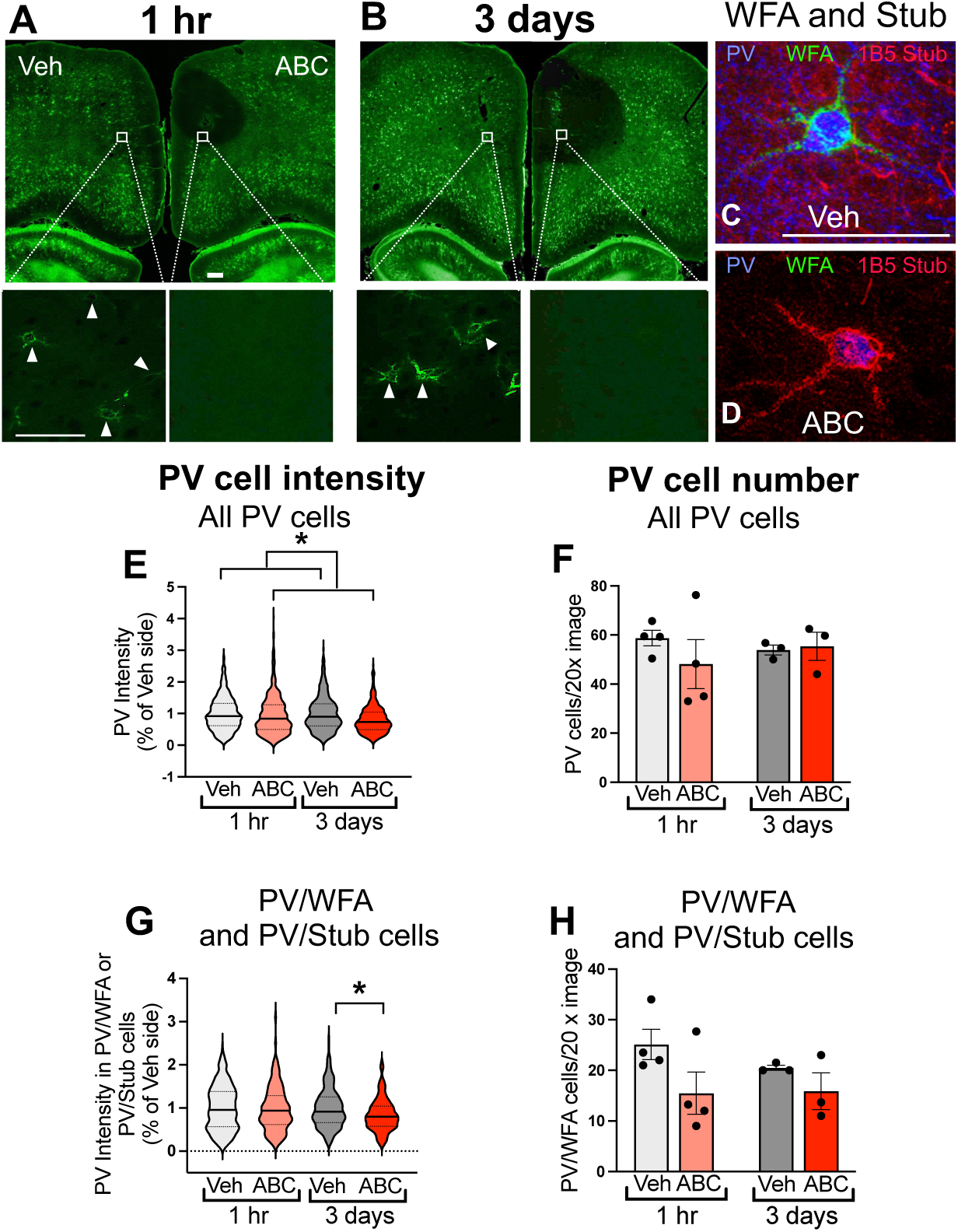
ABC treatment in the mPFC reduces PV cell intensity. **(A, B)** Confocal image of WFA-stained brain (green) showing dorsal PFC area after unilateral Veh or ABC treatment at 1 hr **(A)** or 3 days **(B)** post-injection. Bottom panels show a 20x magnification of the white boxed areas in panels **A** and **B**. **(C, D)** Confocal micrographs of single PNN identified by WFA staining (green) surrounding a PV neuron (blue) from Veh side **(C)** and labeling of “stubs” after ABC treatment identified with 1B5 Stub antibody (red) **(D).** Scale bars for panels **A, B** = 100 µm and **C, D** = 50 µm. **(E)** Intensity of all PV cells 1 hr or 3 days after Veh or ABC treatment. A two-way ANOVA showed a main effect of treatment (F_1,2420_ = 36.78, p < 0.0001), with decreases in PV intensity after ABC treatment. Number of rats (number of cells) for each group: 1 hr: Veh N = 4 (746); ABC N = 4 (546); 3 days: Veh N = 3 (541); ABC N = 3 (561). **(F)** Number of all PV cells was not different after ABC treatment. Number of rats for each group: 1 hr: Veh N = 4; ABC N = 4; 3 days: Veh N = 3; ABC N = 3. **(G)** Intensity of PV/WFA or PV/Stub double-labeled cells. A two-way ANOVA showed a main effect of treatment (F_1,826_ = 4.171, p = 0.0414) and a treatment x time interaction (F_1,826_ = 4.653, p = 0.0313). At 3 days after ABC treatment, PV intensity in PV/WFA double-labeled cells was reduced (p = 0.0082). Number of rats (number of cells) for each group: 1 hr: Veh N = 4 (296); ABC N = 4 (206); 3 days: Veh N = 3 (162); ABC N = 3 (166). **(H)** Number of PV/WFA or PV/Stub double-labeled cells was not different across days or treatment. Number of rats for each group: 1 hr: Veh N = 4; ABC N = 4; 3 days: Veh N = 3; ABC N = 3. For intensity, violin plots show the median and quartiles; for cell number, data are mean ± SEM. * p < 0.05.

We also determined whether ABC treatment altered excitatory or inhibitory puncta surrounding PV cells (***Supplemental Figure 2***). In a subset of the animals used for analysis of PV intensity and cell number (above), we quantified the number of GAD 65/67 and vGLUT1-labeled puncta apposing PV neurons on the Veh-treated side and the ABC-treated side. Note that puncta were assessed around PV neurons with WFA in the case of Veh treatment and PV neurons without WFA in the case of ABC treatment; thus, a subset of PV neurons in the latter group may not have been originally surrounded by PNNs. There was a main effect of time of treatment, with reductions in the number of GAD 65/67 puncta on both Veh- and ABC-injected hemispheres 3 days after injection compared with the number of puncta assessed 1 hr after injection (***Supplemental Figure 2C***). Similarly, the number of vGLUT1 puncta was reduced in both Veh and ABC hemispheres at 3 days vs. at 1 hr following Veh or ABC treatment (***Supplemental Figure 2D***). The ratio of GAD65/67 to vGLUT1 apposing PV neurons was increased in both groups at 3 days, with greater variability (***Supplemental Figure 2E; Supplemental Table 7***). These findings suggest that the microinjection procedure itself, regardless of injectate, had a global effect on the number of puncta surrounding PV neurons with a reduction at 3 days post-injection as compared to 1 hr post injection.

### ABC treatment in the mPFC alters intrinsic properties of FSIs

To examine the impact of degrading PNNs on neuronal excitability and intrinsic properties of FSIs, we employed whole-cell electrophysiology techniques on brain slices from naive rats (see also ***Supplemental Table 8***). ***Figure 4A*** shows the timeline for assessing the intrinsic properties of FSIs (which contain PV) ^54^ within layer V of the prelimbic mPFC at 1 hr or 3 days following *in vivo* Veh or ABC treatment in the mPFC. Several properties were evaluated, including capacitance, input resistance, resting membrane potential (RMP), action potential (AP) half-width, AP threshold, AP amplitude, afterhyperpolarization amplitude (AHP), first interspike interval (ISI), and rheobase (***Figure 4B***). As expected, neuronal excitability in FSIs increased as current steps increased (***Figure 4C***), and there was a current step x time interaction, but only a trend for an interaction between treatment and current injection on neuronal excitability (p = 0.0995). ***Figure 4D*** shows representative traces at 500 pA for the four experimental groups. We found no differences in firing variability among groups (***Figures 4E and 4F***). ABC treatment increased capacitance at both time points (***Figure 4G***), consistent with a previous report ^65^. In addition, the timing of slice preparation after surgery affected some FSI intrinsic properties. There was higher input resistance at 3 days compared with 1 hr after both Veh and ABC treatment (***Figure 4H***), an increase in action potential amplitude for both groups at 3 days (***Figure 4L***), and a decrease in rheobase current at 3 days (***Figure 4O***), together suggesting that there may have been damage-induced decreases in some of these properties 1 hr after disturbance of tissue by microinjection or alternatively, that inflammatory responses produced an increase in 3 days later. There were no changes in resting membrane potential (***Figure 4I***) or action potential half-width (***Figure 4J***), but ABC treatment decreased the action potential threshold at both 1 hr and 3 days (***Figure 4K***), suggesting that PNNs may alter the response of PV neurons to incoming signals. We found no increases in afterhyperpolarization amplitude (***Figure 4M***), but there was an increase in the first inter-spike interval at 1 hr, but not 3 days, after ABC treatment (***Figure 4N***), suggesting that PNN removal delays firing.

**Figure 4.**
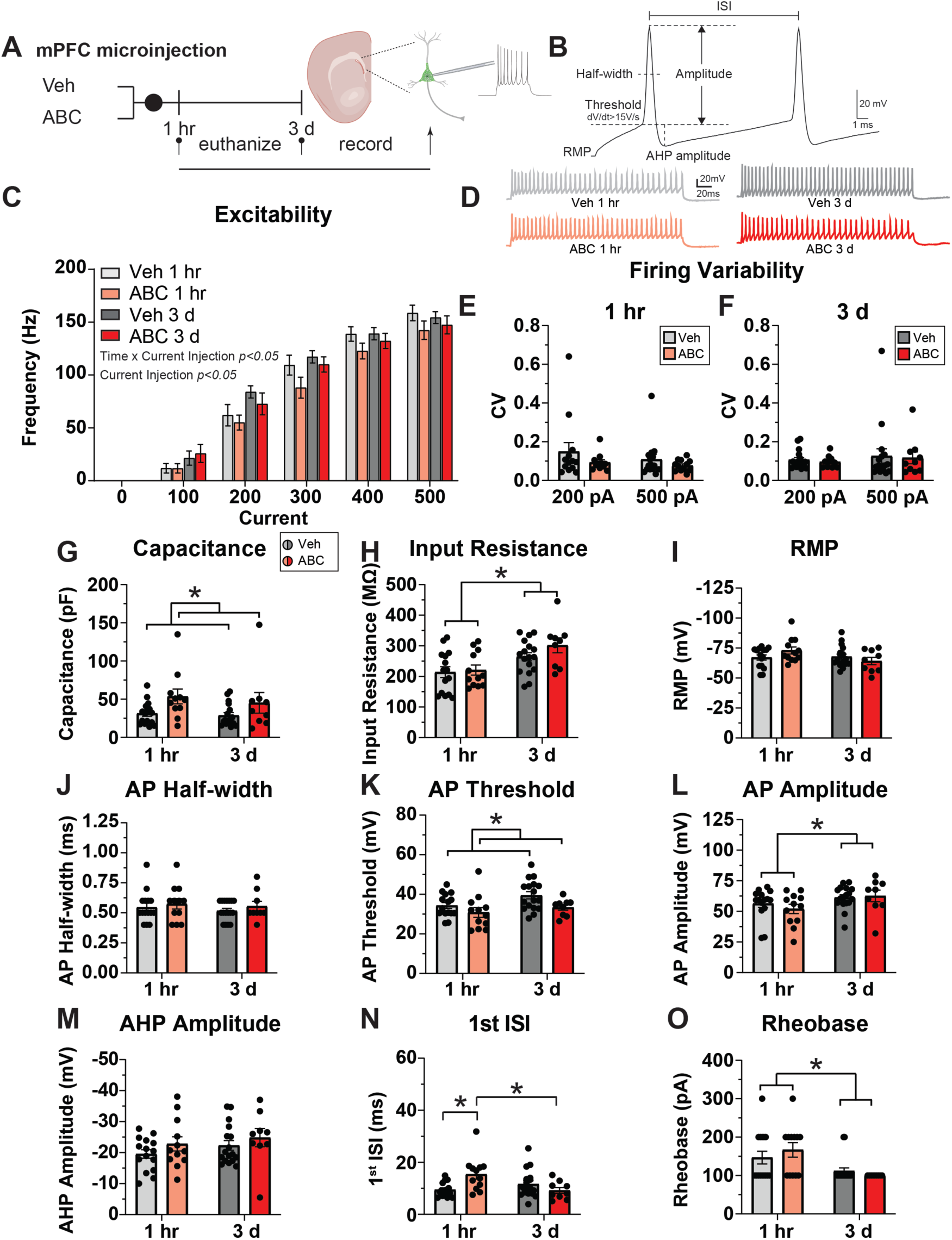
ABC treatment in the mPFC alters intrinsic properties of FSIs. **(A)** Timeline of treatment. **(B)** Description of parameters. **(C)** PNN removal attenuated neuronal excitability as current steps increased. A three-way RM ANOVA showed a main effect of current step (F_2.819,138.1_ = 628.7, p < 0.0001) and a time x current step interaction (F_6,294_ = 2.565, p = 0.0195). Number of rats (number of cells): 1 hr: Veh N = 7 (15); ABC N = 4 (12); 3 days: Veh N = 10 (17); ABC N = 7 (9). **(D)** Representative traces at 500 pA for the four experimental groups. **(E)** There were no differences in firing variability at 1 hr in either treatment group. Number of rats (number of cells): 200 pA: Veh N = 7 (13); ABC N = 4 (11); 500 pA: Veh N = 7 (15); ABC N = 4 (12). **(F)** There were no differences in firing variability at 3 days in either treatment group. Number of rats (number of cells): 200 pA: Veh N = 10 (17); ABC N = 7 (10); 500 pA: Veh N = 10 (17); ABC N = 7 (10). **(G)** PNN removal increased membrane capacitance at both 1 hr and 3 days. A two-way ANOVA showed a main effect of treatment (F_1,49_ = 7.415, p = 0.0089). **(H)** Input resistance was increased at 3 days vs. 1 hr; a two-way ANOVA showed a main effect of time (F_1,49_ = 13.73, p = 0.0005). **(I)** There were no differences in resting membrane potential (RMP) at 1 hr or 3 days in either treatment group. **(J)** There were no differences in the AP half-width at 1 hr or 3 days in either treatment group. **(K)** PNN removal reduced the AP threshold at both 1 hr and 3 days, with a main effect of treatment (F_1,49_ = 5.984, p = 0.0181). **(L)** The AP amplitude was increased at 3 days vs. 1 hr in both groups, with a main effect of time (F_1,49_ = 5.170, p = 0.0274). **(M)** There were no differences in the afterhyperpolarization (AHP) amplitude at 1 hr or 3 days in either treatment group. **(N)** PNN removal increased the first interspike interval time at 1 hr, but not at 3 days. A two-way ANOVA showed a time x treatment interaction (F_1,49_ = 9.689, p = 0.0031), with post-hoc analysis showing an effect of ABC at 1 hr (Veh vs ABC: p = 0.0046) and an effect across time in ABC-treated rats (p = 0.0093). **(O)** Rheobase current was reduced at 3 days vs. 1 hr, with a main effect of time (F_1,49_ = 12. 99, p = 0.0007). Data are mean ± SEM. For **(H – 0)** number of rats (number of cells): 1 hr: Veh N = 7 (15); ABC N = 4 (12); 3 days: Veh N = 10 (17); ABC N = 7 (9). * p < 0.05.

### ABC treatment does not alter GluN2B receptors in the mPFC after novel VR5 reactivation session

The blockade of GluN2B receptors prevents the ability to render labile a reactivated memory for disruption by amnestic agents ^66–68^. More recent evidence suggests that GluN2B receptors may only play a role in destabilizing more recent memories and are not the mechanism that may support novelty induced destabilization ^69^. Nevertheless, we measured the level of surface and intracellular GluN2B subunits in both the mPFC and amygdala ^66^ 30 min after FR1 or VR5 memory reactivation (***Supplemental Figures 3A-E; Supplemental Table 9***). We found no differences in surface or intracellular GluN2B subunits in either brain area, nor did we find differences between treatment groups in the ratio of surface:intracellular GluN2B levels (data not shown). GluN2B receptor subunit trafficking may therefore not be sensitive to the differences in FR1 vs. VR5 memory reactivation, or the changes may be too small to measure when assessing whole tissue punches.

### Increased mPFC beta and gamma power after novel VR5 reactivation session is prevented by ABC in the mPFC during rewarded lever presses

To determine the extent to which the novel VR5 reactivation session altered LFPs within the mPFC or coupling with the hippocampus and whether ABC would modify this coupling, we measured LFPs within the mPFC and dHIP on the last day of self-administration training and during the VR5 reactivation session in a subset of animals. All animals were given ABC 3 days prior to the VR5 reactivation as in ***Figure 2A*** (Pre-VR5 group). ***Figure 5A*** shows the raw and filtered traces for LFPs in the mPFC. Spectrograms from the mPFC show that rewarded lever presses on the novel VR5 session compared with the familiar last day of self-administration training were accompanied by an increase in both 45-65 Hz gamma and 35-45 Hz gamma power (***Figures 5B, 5C upper panels, 5E, 5F***) and also in 15-30 Hz (beta/low gamma) power (***Figures 5B, 5C upper panels, 5G***) following rewarded lever presses. For rewarded lever presses, the increase in gamma power (35-45 H) after novel VR5 reactivation was prevented by ABC treatment in the mPFC (***Figures 5C, 5D upper panels, 5I***), with a strong trend toward decreased beta/low gamma power as well after ABC (***Figures 5C, 5D upper panels, 5J***). In addition, during the novel VR5 reactivation session, gamma power was increased in rewarded compared with unrewarded lever presses, and this difference was not observed after ABC treatment (***Figures 5C, 5D upper panels, 5H, 5I***). The visually apparent increase in delta power (2-4 Hz) from Veh-treated animals after VR5 reactivation (***Figure 5C, lower panel)*** was not significantly different among any groups. Thus, a novel VR5 reactivation elevated beta and gamma activity in response to rewarded lever presses compared with the familiar last day of training, and PNN removal prevented these elevated responses on the VR5 reactivation session. (Statistical data shown in ***Supplemental Table 10***).

**Figure 5.**
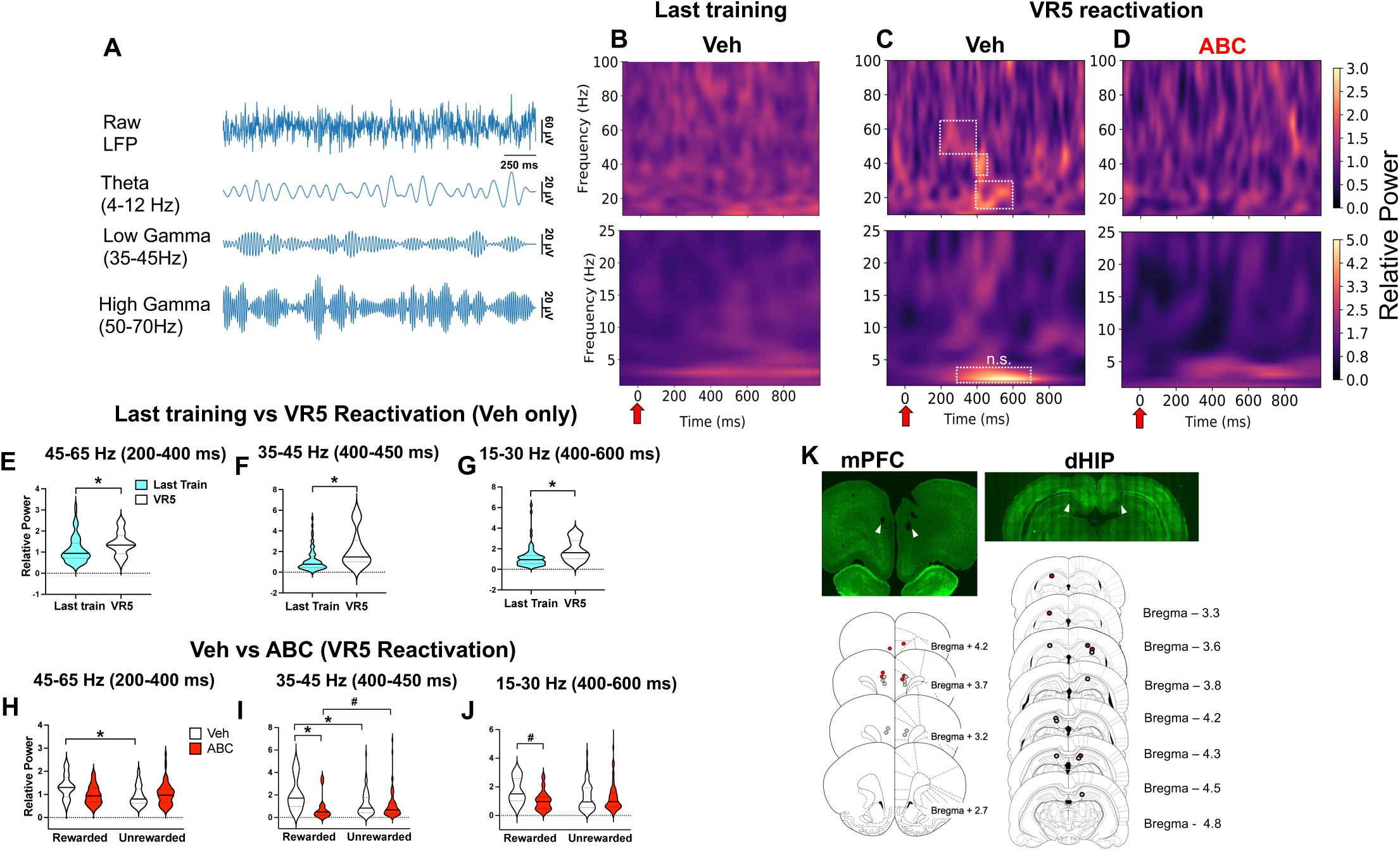
Novel VR5 memory reactivation session: ABC treatment in the mPFC prevents increases in beta/gamma power during rewarded lever presses. **(A)** Raw and filtered LFP traces from mPFC. **(B)** Spectrograms of mPFC power in the 10-100 Hz range (upper panel) and 0-25 Hz range (lower panel) after rewarded lever presses on the last training day. **(C)** Spectrograms of mPFC power in Veh rats in the 10-100 Hz range (upper panel) and 0-25 Hz range (lower panel) after rewarded lever presses on the VR5 reactivation session. **(D)** Spectrograms of mPFC power in ABC rats in the 10-100 Hz range (upper panel) and 0-25 Hz range (lower panel) after rewarded lever presses on the VR5 reactivation session. For B-D: color bar is relative power. **(E)** Quantification of 45-65 Hz power (upper white square in 5C, upper panel) in the mPFC 200-400 ms after rewarded lever presses during last day training vs. VR5 reactivation session in Veh rats. There were no differences in this gamma range comparing across the two days. Note that ABC was not delivered until after the last training session, and therefore only Veh animals are shown for the last training day for direct comparison with the Veh group on the VR5 reactivation session. **(F)** Quantification of 35-45 Hz power (middle white rectangle in 5C, upper panel) in the mPFC 400-450 ms after rewarded lever presses showed an increase in power during the VR5 session vs the last training day (p = 0.0018). **(G)** Quantification of 15-30 Hz power (lower white rectangle in 5C, upper panel) in the mPFC 400-600 ms after rewarded lever presses showed an increased power during the VR5 session vs the last training day (p = 0.0287). There were no differences in the 2-4 Hz range at 300-700 ms after rewarded lever presses among groups (*Supplemental Table 10*). **(H)** Quantification of 45-65 Hz power (upper white square in 5C, upper panel) in the mPFC 200-400 ms after rewarded and unrewarded lever presses. A two-way ANOVA showed a main effect of lever (F_1, 170_ = 4.142, p = 0.0434) and a treatment x lever interaction (F_1, 170_ = 5.271, p = 0.0229). In Veh rats, post-hoc analysis showed a decrease in power in the unrewarded vs rewarded lever presses (p = 0.0058). **(I)** Quantification of 35-45 Hz power (middle white rectangle in 5C, upper panel) in the mPFC 400-450 ms after rewarded and unrewarded lever presses. A two-way ANOVA showed a main effect of Veh vs ABC treatment (F_1, 170_ = 16.67, p < 0.0001) and a treatment x lever interaction (F_1, 170_ = 11.37, p = 0.0009). Post-hoc analysis revealed that rewarded lever presses were accompanied by increased power in Veh vs ABC treated rats (p < 0.0001) and in rewarded vs unrewarded lever presses in Veh rats (p = 0.0232), with a trend toward decreased power in rewarded vs unrewarded lever presses in ABC rats (p = 0.0552). **(J)** Quantification of 15-30 Hz power (lower white rectangle in 5C, upper panel) in the mPFC 400-600 ms after rewarded and unrewarded lever presses. A two-way ANOVA showed a treatment x lever interaction (F_1, 170_ = 4.129, p = 0.0437), with a strong trend toward reduced power in the rewarded lever presses between Veh and ABC treated rats (p = 0.0556). **(K)** Location of electrodes in a representative mPFC (top left panel) and dHIP (top right panel). Location of all electrodes in the mPFC (left bottom panel) and location of all electrodes in the hippocampus, with exception of one electrode site that was not identified in one ABC rat (right bottom panel). Gray dots = Veh; red dots = ABC. Number of rats (number of trials): Veh N = 5 (16 rewarded, 70 unrewarded); ABC N = 3 (17 rewarded, 71 unrewarded). *p < 0.05; # p < 0.06.

### Increased dHIP/mPFC coupling during novel VR5 reactivation session is reduced by ABC treatment in the mPFC

Given that PV neurons coordinate long-range communication within and across other brain regions through coupling of theta and gamma oscillations ^70^, we also examined cross-frequency phase-amplitude coupling (PAC) between dHIP theta phase and mPFC gamma amplitude. Comodulograms 200-600 ms after rewarded lever presses are shown for rats during the last day of training given Veh (***Figure 6A***), for Veh rats during the VR5 reactivation session (***Figure 6B***), and for ABC rats during the VR5 reactivation session (***Figure 6C***). Coupling was apparent between dHIP 8-10 Hz theta phase and mPFC 50-70 Hz gamma frequency in the Veh group during the VR5 reactivation session. This same coupling was absent on the last day of training in the same Veh animals (***Figure 6A***). In addition, on the VR5 reactivation session, ABC reduced this coupling (***Figure 6C***). Based on the comodulograms, we tested for changes in the distribution of gamma power (50-70 Hz) over theta phase (8-10 Hz). There was a significant dHIP theta phase modulation of mPFC gamma power 200-600 ms after rewarded lever presses in Veh rats during the VR5 session (***Figure 6E***). This modulation was absent in the same Veh rats during the last training day (***Figure 6D***). During the VR5 reactivation, ABC treatment appeared to reduce, but not completely prevent, dHIP modulation of mPFC compared with Veh controls (***Figure 6F***). Thus, the novel VR5 reactivation session was associated with coupling between the dHIP and mPFC, and PNN removal dampened this coupling. The reduced coupling in ABC-treated rats was not likely to be due to differences in lever-pressing activity in general, since the two groups did not differ in lever pressing during the VR5 reactivation session (see Figure 2D).

**Figure 6.**
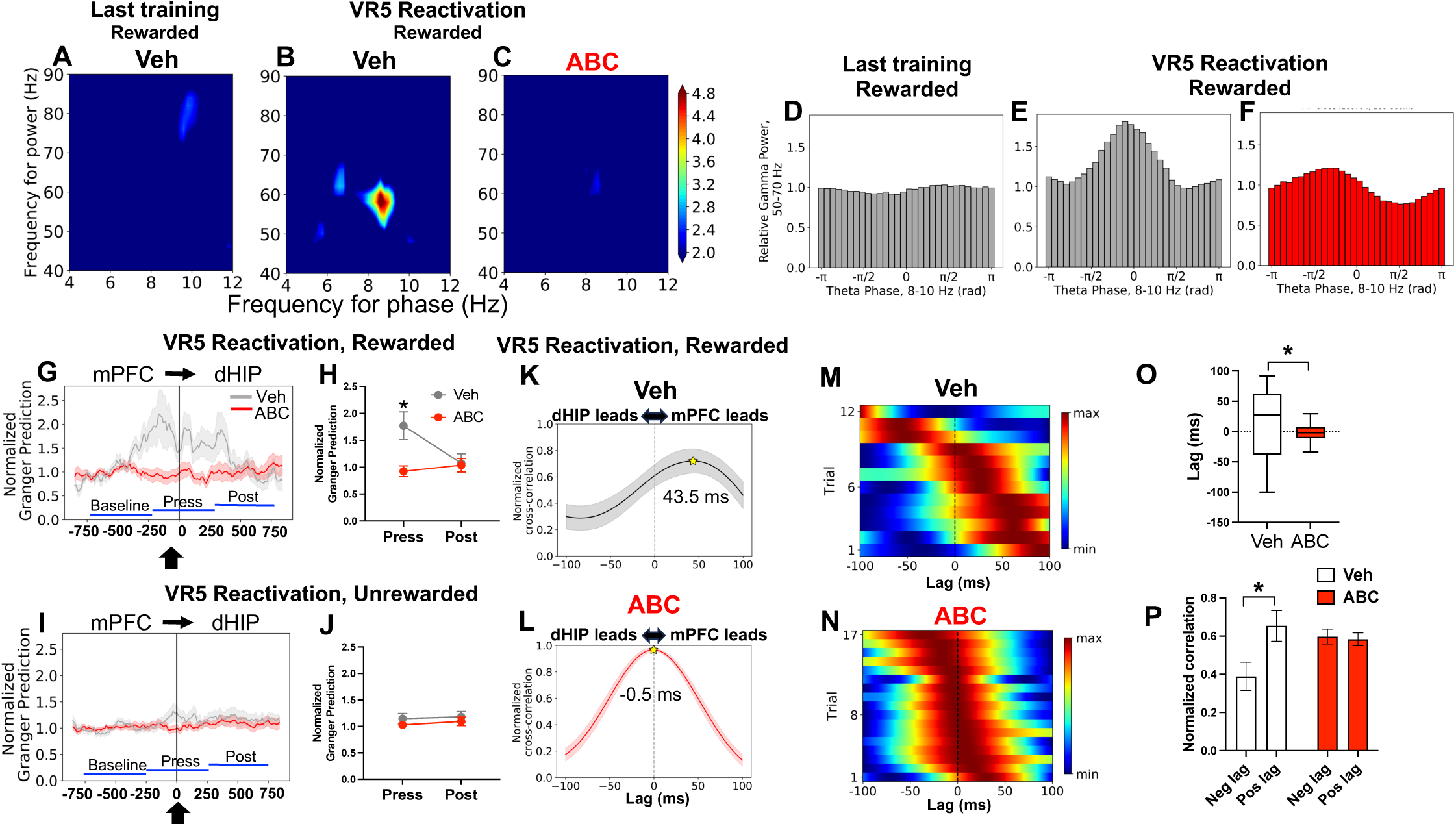
Increased dHIP/mPFC coupling and directional flow between the mPFC and dHIP during novel VR5 memory reactivation is prevented by ABC treatment in the mPFC. **(A)** Veh rats on last training day: comodulogram showing dHIP theta phase and mPFC amplitude 200-600 ms after rewarded lever presses. **(B)** Veh rats on VR5 reactivation session: comodulogram showing dHIP theta phase and mPFC amplitude 200-600 ms after rewarded lever presses. Note increased coupling of dHIP theta in the 8-10 Hz range with mPFC gamma in the 50-70 Hz range. **(C)** ABC rats on VR5 reactivation session: comodulogram showing dHIP theta phase and mPFC amplitude 200-600 ms after rewarded lever presses. Note absence of coupling observed in Veh rats. Color bar is z-score; comodulograms are floored at p < 0.05. **(D)** Veh rats on last training day: plot of mPFC gamma power (50-70 Hz) binned by dHIP theta phase (8-10 Hz) 200-600 ms after rewarded lever presses. There was no significant modulation of mPFC gamma power by dHIP theta (Modulation index (MI) = 1.92 x 10^-4^; z = −0.77). **(E)** Veh rats on VR5 reactivation session: plot of mPFC gamma power (50-70 Hz) binned by dHIP theta phase (8-10 Hz) 200-600 ms after rewarded lever presses. dHIP theta significantly modulates mPFC gamma power after rewarded lever presses (MI = 6.59 x 10^-3^; z = 2.12). **(F)** ABC rats on VR5 reactivation session: plot of mPFC gamma power (50-70 Hz) binned by dHIP theta phase (8-10 Hz) 200-600 ms after rewarded lever presses. dHIP theta significantly modulates mPFC gamma power (MI = 3.43 x 10^-3^; z = 1.89). **(G)** Granger causality (GC) analysis computed from the raw LFP signal in the dHIP and mPFC in 500 ms bins for rewarded lever presses prior to lever press (−750 to −250 ms, baseline), surrounding lever press (−250 to 250 ms), and after lever press (250 to 750 ms) during VR5 memory reactivation session. **(H)** GC normalized to its own averaged baseline for Veh or ABC groups during rewarded lever press and post-lever press. Two-way RM ANOVA shows a main effect of treatment (F_1, 27_ = 4.892, p = 0.0356), time (F_1, 27_ = 6.818, p = 0.0146), and a treatment x time interaction (F_1, 27_ = 13.20, p = 0.0012). The mPFC leads the dHIP around rewarded lever presses but disappears by 250-750 ms after lever press. **(I)** GC analysis as in **(G)** for unrewarded lever presses. **(J)** Normalized GC to its own averaged baseline showing no changes between Veh and ABC groups. **(K, L)** Cross-correlation between mPFC/dHIP theta (4-12 Hz) amplitude envelopes at varying lags in Veh **(K)** and ABC **(L)** groups. Stars show peak correlation (mean ± SEM: Veh = 43.5 ms ± 0.03 ms; ABC = −0.5 ± 0.003 ms). Line is mean, shading represents SEM. **(M, N)** Normalized heat maps of theta amplitude cross-correlation by trial from rewarded trials in Veh- and ABC-treated rats in Veh **(M)** and ABC **(N)** groups. Darker red color indicates higher cross-correlation. **(O)** Lag at peak correlation, with a significant difference in distributions between the Veh and ABC groups (Komogorov-Smirnov p = 0.0111). Box plot shows median and 95% confidence intervals. **(P)** Normalized correlation for negative (Neg) and positive (Pos) lag, with positive lag indicating mPFC leading the dHIP. There was a main effect of lag (F_1,54_ = 5.241; p = 0.0260) and a significant treatment x lag interaction (F_1,54_ = 6.477; p = 0.0138). Post-hoc analysis showed a significant lead by the mPFC in the Veh (p = 0.0052) but not in the ABC group. Number of rats (number of trials): Veh N = 4 (12 rewarded, 59 unrewarded); ABC N = 3 (17 rewarded, 71 unrewarded). *p < 0.05.

Functional interaction between the dHIP and mPFC was assessed using Granger Causality (GC) analysis to measure the direction and extent to which information flowed between the mPFC and dHIP during rewarded and unrewarded lever presses on the VR5 session (***Figures 6G-J***). GC analysis suggested a significant flow of information from the mPFC to the dHIP during rewarded lever presses in the Veh but not in the ABC group (***Figures 6G, 6H***), with no differences during unrewarded lever presses (***Figures 6I, 6J***). This mPFC lead appeared just prior to the rewarded lever press and disappeared within 250 ms after the lever press. To confirm these findings, we computed cross-correlation between mPFC/dHIP theta (4-12 Hz) amplitude envelopes at varying lags in Veh and ABC groups (***Figures 6K, 6L***). ***Figures 6M, 6N*** show normalized heat maps of theta amplitude cross-correlations by trial from rewarded trials in Veh- and ABC-treated rats, and ***Figure 6O*** shows the distribution of lag at peak correlation, which was different between ABC- and Veh-treated rats. ***Figure 6P*** shows a higher cross correlation for the positive lag (mPFC lead) compared with the negative lag (dHIP lead), with no differences in the ABC group. Thus, consistent with GC analysis, ABC disrupted the ability of the mPFC to lead the dHIP during rewarded lever presses. These findings suggest that PNNs influence directional connectivity from the mPFC to dHIP during cocaine-seeking/taking behavior. All statistical data are shown in ***Supplemental Table 10***.

We also assessed dHIP theta/mPFC gamma coupling during *unrewarded* lever presses. Unlike for rewarded lever presses, there was no significant modulation of mPFC 50-70 Hz gamma by dHIP 8-10 Hz theta in either the Veh or ABC group (***Supplemental Figures 4A-D***). Lag analysis of mPFC/dHIP theta amplitude cross-correlation indicated that there was a significant difference in distributions of peak theta lag between the Veh and ABC groups (***Supplemental Figures 4H***). There was a main effect of treatment for the normalized correlation coefficient, which was lower in the Veh- compared with ABC-treated groups (***Supplemental Figures 4E - 4G***), but there was no significant lead by the mPFC or dHIP in either group. Thus, unlike for rewarded lever presses, we did not see dHIP/mPFC coupling or directional flow from the mPFC to dHIP for unrewarded lever presses. All statistical data for Supplemental Figure 4 are shown in ***Supplemental Table 11***.

### ABC treatment alters inter-hemispheric mPFC theta/gamma coupling after unrewarded lever presses during novel VR5 reactivation session

Given that PV interneurons support cross-hemispheric coupling via direct callosal projections ^71^, we also examined the effect of ABC treatment on inter-hemispheric theta gamma coupling across the mPFC. The most apparent differences in coupling were found after unrewarded lever presses. During the VR5 reactivation session, there was significant cross-hemispheric mPFC 8-10 Hz theta phase modulation of mPFC 70-90 Hz gamma power 200-600 ms after unrewarded lever presses in Veh rats during the VR5 session (***Supplemental Figures 4I, 4K***), and ABC blocked modulation in this frequency range (***Supplemental Figures 4J, 4L***). However, we also found significant cross-hemispheric mPFC modulation of gamma by theta in the 5-7 Hz range in the ABC but not in the Veh group (***Supplemental Figures 4M, 4N***). Lag analysis of cross-hemispheric mPFC theta amplitude envelope correlations revealed a main effect of lag, with no effect of ABC treatment (***Supplemental Figures 4O - 4Q***). There were no differences in the distribution of lag at peak correlation between Veh and ABC treated groups (***Supplemental Figure 4R***). These results indicate that unrewarded lever presses may be associated with inter-hemispheric mPFC coupling, with a different theta frequency modulating gamma activity in Veh- vs ABC-treated rats. All statistical data are shown in ***Supplemental Table 11***.

## Discussion

These studies explored the cellular and connectivity mechanisms in mPFC associated with cocaine memory reconsolidation, which is important for understanding how to reduce relapse. We report several new major findings in which we use ABC to remove PNNs in the mPFC: 1) PNN removal in the mPFC prior to training for cocaine self-administration attenuated the acquisition of cocaine self-administration; 2) PNN removal 3 days prior to, but not within ∼ 1 hr after a memory reactivation session blocked cue-induced reinstatement and reduced the reinforcing effects of cocaine. However, the reduction in cue reinstatement was present *only in rats given a novel reactivation session*; 3) PNN removal in naive rats reduced the intensity of PV cells surrounded by PNNs 3 days, but not 1 hr later, with no differences in excitatory or inhibitory puncta surrounding PV neurons; 4) In mPFC slices, PNN removal increased membrane capacitance at both 1 hr and 3 days later and delayed the first interspike interval at 1 hr but not 3 days later; 5) *In vivo* recordings revealed increased high frequency power and increased coupling between dHIP theta phase and mPFC gamma power after rewarded lever presses during a novel reactivation session but not during the familiar last day of self-administration training. These increases in coupling were reduced after PNN removal, and the mPFC directional flow to the dHIP during rewarded lever presses was also reduced after ABC treatment; 6) Theta/gamma coupling across mPFC hemispheres was increased during *unrewarded* lever presses; this coupling was altered after PNN removal. Thus, PNN removal generally reduced or altered communication between the mPFC and dHIP and across the mPFC, which may be necessary for re-stabilizing networks during cocaine memory updating.

### PNN removal in the dorsal mPFC attenuates acquisition of cocaine self-administration, blocks cue-induced reinstatement, and reduces reinforcing efficacy of cocaine

Similar to our previous study using cocaine-induced CPP ^22^, here we found that PNN removal reduced the acquisition of cocaine self-administration in rats. We also showed in CPP studies that removing PNNs with ABC in the mPFC prior to memory reactivation blunted the reconsolidation of a cocaine-induced CPP memory ^22^; however, a self-administration memory entails a much greater number of pairings, approximately 250-300 between the cocaine cue light compared with three pairings in CPP ^22^. Memory reconsolidation is harder to disrupt in well-established and older memories ^72, 73^. Indeed, in the present study, we found no effect of PNN removal in animals trained and given a memory reactivation session under their familiar FR1 schedule. These findings were similar to our previous observation that MK801 given during memory reactivation blocked the reconsolidation of a cocaine CPP memory but not a cocaine self-administration memory ^74^.

Several studies have demonstrated that, to disrupt stronger memories, *novel information* is necessary to destabilize the preexisting memory and allow for memory updating and subsequent reduced recall of the old memory ^27–30^. A likely role for the reconsolidation process that occurs in response to novelty is that it serves to update memories by creating a prediction error between what is expected and what is received ^30, 75–85^. Our current study is based on previous work showing that reconsolidation of well-learned memories for self-administration of sucrose ^27, 29^ and cocaine ^27^ were disrupted by changing the reinforcement schedule during the memory reactivation session so that it is different from the one on which the rats were trained, which requires a change in strategy to obtain the reward (additional lever presses). A novel or unpredictable component that is delivered alongside the unconditioned stimulus (cocaine in our study) may be a key element in updating strong memories ^86–89^. Park et al. ^90^ showed that exposure to novelty resulted in the initial decoupling of the ventral hippocampus from the mPFC and subsequent re-potentiation of this circuit. PNN removal may similarly prevent the recoupling of circuits linking the dHIP to the mPFC. In support of these results, we found that the novel VR5 reactivation session increased coupling between dHIP theta phase and mPFC gamma amplitude that was absent on the last day of self-administration, when the memory was familiar. Thus, coupling between these brain regions may be a marker for the circuit impact of novelty and subsequently destabilizing and updating cocaine-associated memories.

The lack of effect of PNN depletion on responding during the memory reactivation session itself suggests that PNN removal does not rely on the mPFC circuit during the retrieval process. Alternatively, it suggests that cocaine-induced increases in firing of mPFC neurons ^91, 92^, including PV neurons ^93^, may have overridden the impact of PNN removal on the retrieval process. The effect of ABC and VR5 reactivation persisted under a PR schedule of reinforcement in which cocaine was available, supporting the idea that PNN removal after a novel VR5 reactivation reduces the reinforcing efficacy of cocaine. Our findings are in agreement with those from a recent study in which PNN removal in the entorhinal cortex disrupted the grid cell network for a spatial memory, but when animals were placed into a *novel environment* to form a new memory, PNN removal degraded spatial representation of both the new and old memories ^31^. Similar to their findings and those from others^18, 39^, we observed that PNN removal in the mPFC did not cause apparent changes in network function (as measured by behavior) until the system was required to process new information. As long as the schedule of reinforcement was familiar, PNN removal did not disrupt the representation of the old memory. The lack of an effect of PNN removal in animals self-administering sucrose is consistent with the idea that cocaine induces a state of metaplasticity^94^ that sucrose does not. Several studies show that treatments that block reinstatement of cocaine seeking do not block sucrose seeking ^95–97^, with different neural ensembles mediating food or sucrose vs. those mediating cocaine seeking ^42, 98, 99^.

### mPFC contributes to memory reconsolidation: possible role of extinction

We gave a 30 min extinction session prior to cue-induced reinstatement to assess 1) whether extinction responding was altered by PNN removal, and 2) the degree to which contingent presentation of the cue produced reinstatement. This extinction session may have influenced subsequent reinstatement and PR responding after PNN removal. We previously found in cocaine self-administering rats that microinjection of anisomycin into the mPFC to inhibit protein synthesis prior to memory reactivation suppressed subsequent reinstatement by cocaine + cue, but only in rats that also received extinction prior to memory reactivation ^100^. Thus, extinction may have recruited mPFC circuitry. Our findings that PNN removal prior to but not after memory retrieval reduced reinstatement are similar to those from a fear conditioning study ^101^. In that study, PNN removal given *prior to* fear training prevented the re-emergence of fear by renewal or spontaneous recovery, but PNN removal *after* training had no effect on re-emergence of fear expression after extinction ^101^. Thus, retrieval under novel conditions may create instability in the mPFC circuit required for reconsolidation, which may be dependent on intact mPFC PNNs *at the time of memory reactivation*. The absence of an effect when animals were given no memory reactivation or the same, familiar reactivation as that during training (FR1 and FR3 groups) strongly suggest that reconsolidation of cocaine memory is dependent on intact mPFC PNNs at the time of memory reactivation. Whether PNN removal prior to memory reactivation renders the memory more malleable to extinction-induced processes similar to that described by Gogolla et al. ^101^ or instead manifests as a reduced ability to promote memory updating remains to be tested.

### PNN removal alters intrinsic properties of FSIs but does not alter excitatory or inhibitory puncta surrounding FSIs

The increased capacitance of FSIs at both the 1-hr and 3-day timepoints after PNN removal aligns with previous research ^65, 102^. While we did not observe a significant attenuation in AP frequency as previously reported after ABC ^31, 65, 103–105^, we did find a trend towards a reduction in AP frequency, somewhat consistent with modeling studies demonstrating a reduction in firing frequency when capacitance is increased ^102^, the latter which we observed after PNN removal. However, several studies have also reported no effect of ABC on AP frequency ^40, 106–108^; the differences with the current finding may be due to species variability ^104^, variation in ABC treatment protocols and brain areas, and the duration following ABC treatment. The increase in the ISI at the 1-hour timepoint after PNN removal may have altered the precision of feedforward inhibition on pyramidal neuron output. Our findings strengthen the notion that PNNs play a role in shaping AP dynamics. Notably, we assessed PV neuron function after PNN removal in naive rats, in the absence of any history of cocaine self-administration. We previously reported that intra-mPFC injection of ABC led to profound suppression of AP frequency in FSIs 2 hr after cocaine reinstatement in rats trained for cocaine CPP ^63^ and thus, memory reactivation in the presence of cocaine in the current study may have reduced PV neuron firing. Decreased PV intensity levels at 3 days but not 1 day after PNN removal suggests the possibility of suboptimal functioning due to imprecise firing of these neurons. As PV neurons and gamma oscillations critically contribute to mPFC output during rule switching ^71, 109, 110^, their inability to optimally function may have compromised the updating of the cocaine memory.

We previously found that limited cocaine exposure altered inputs to PV neurons in mPFC, including the excitatory:inhibitory ratio^49, 53^. However, here we found that PNN removal did not alter excitatory or inhibitory puncta around PV neurons compared with Veh injections, but both groups showed changes with regard to time after microinjection. These findings suggest that the microinjection itself may have altered the number of excitatory and inhibitory puncta over time, possibly due to physical disruptions from local microinjection.

### Interregional coherence and reconsolidation

Several studies have shown coupling between the dHIP and mPFC in a variety of tasks that involve spatial or working memory ^111–115^. In the current study, strong coupling between dHIP theta and mPFC gamma did not appear necessary for the retrieval of memory, since PNN removal decreased coupling yet demonstrated no difference from Veh controls in active lever pressing during the VR5 memory reactivation session. Coupling between dHIP/mPFC may instead be associated with the destabilization of memory^90^. Our data support the idea that Veh controls were able to process the VR5 session as novel, whereas rats given ABC likely did not: Veh animals showed elevated beta and gamma activity during rewarded lever presses during the VR5 session, and elevated beta activity has previously been described for transient responses to novelty ^116^. The directionality from the mPFC to dHIP during rewarded lever presses was also lost after PNN removal. In addition, PNN removal altered theta/gamma coupling across mPFC hemispheres during *unrewarded* lever presses, with a different theta frequency modulating gamma amplitude in Veh- vs ABC-treated rats. Overall, our findings with the novel memory reactivation session suggest that increased synchrony between the mPFC and dHIP during rewarded lever presses and increased specific theta frequency modulation of gamma across mPFC hemispheres during unrewarded lever presses may serve as error signals, which are reduced in ABC-treated rats.

### Limitations

ABC degrades chondroitin sulfate glycosaminoglycans on chondroitin sulfate proteoglycans ^117^ but also degrades the loose ECM ^118^. While we cannot rule out potential effects of loose ECM removal in our studies, there is evidence that PNNs are a major contributor to altered plasticity after ABC treatment because knockout mice with depletion of PNN components still show enhanced plasticity states ^119, 120^, and additional ABC treatment in knockout mice lacking the gene for the PNN-specific protein Crtl1 showed no further changes in object recognition memory ^121^. An additional limitation is that we used only male rats. Acquisition and reinstatement in cocaine self-administering rats is higher in females ^122, 123^, and our group and others have reported sex differences in PNN intensity in reward-associated brain regions ^124–128^. It is therefore possible that ABC would impact males and females differently, and future studies would also need to take into account the estrous phase ^129, 130^.

### Conclusions

Our findings strongly suggest that removal of PNNs in the mPFC with ABC prevents updating of a cocaine-associated memory, but only when the reactivated memory is destabilized by introducing a novel component to the experience. If PNNs are removed after memory reactivation, reconsolidation is still successful. The novel reactivation session increases beta and gamma activity in the mPFC, and disrupted synchrony between dHIP theta phase and mPFC gamma power may prevent memory updating in animals lacking PNNs in the mPFC. Collectively, our findings support the idea that a novel component presented during cocaine memory reactivation increases both beta/gamma activity in the mPFC and coupling between the dHIP and mPFC, and that a PNN-dependent stable network is needed for updating of that memory.

## Acknowledgements

The authors thank Ryan Todd, Drs. Jordan Blacktop, Megan Slaker, and Lynn Churchill for assistance with the acquisition studies.

## Funding

This work was supported by DA 040965 (BAS, TEB, SAA), DA 055645 (BAS, TEB, SAA, AIA), OHSU Physician Scientist Award, Portland Veterans Affairs Research Foundation (AIA), and the Good Samaritan Foundation of Legacy Health (BAS).

---

**Supplemental Figure 1.**
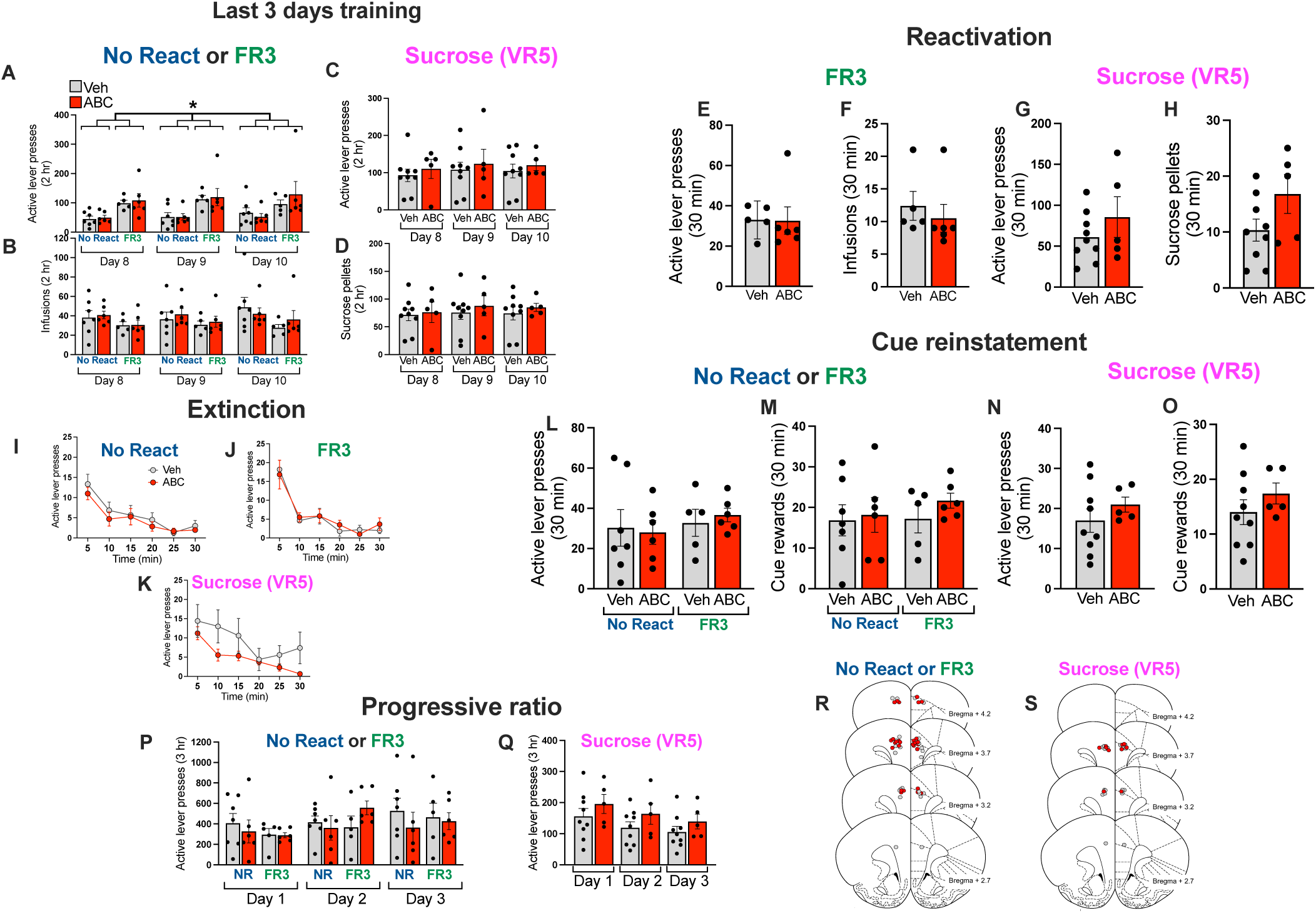
ABC treatment in the mPFC does not block cocaine cue-induced reinstatement in the absence of memory reactivation, nor does it block sucrose cue-induced reinstatement after a novel VR5 memory reactivation session. **(A)** Active lever presses during the last 3 days of training in rats trained on an FR1 schedule (No React and Sucrose groups) or an FR3 schedule (FR3) as in Figure 2. A three-way RM ANOVA showed a main effect of reactivation type (F_1,20_ = 9.426, p = 0.0060), with increased lever pressing in the FR3 group vs. No React groups (trained on an FR1). **(B)** Number of cocaine infusions was not different across the last 3 days of training. **(C)** Number of active lever presses for sucrose pellets was not different across the last 3 days of training (trained on an FR1, reactivated on a VR5). **(D)** Number of sucrose rewards was not different across the last 3 days of training. **(E-H)** During memory reactivation, rats trained on the FR3 schedule were also given an FR3 schedule for memory reactivation, while rats trained for sucrose pellets were given a VR5 schedule for memory reactivation. In the FR3 groups, active lever presses **(E)** or cocaine infusions **(F)** were not different between Veh and ABC groups, and in the Sucrose VR5 groups, active lever presses **(G)** or number of sucrose pellets **(H)** were not different between Veh and ABC groups. **(I-K)** Active lever presses during extinction in No React **(I)**, FR3 **(J)**, and Sucrose VR5 **(K)** groups. A two-way RM ANOVA showed only an effect of time in all three groups (No reactivation: F_2.426, 26.69_ = 14.01, p< 0.0001; FR3: F_2.085, 18.77_ = 21.25, p < 0.0001; sucrose F_3.630, 43.56_ = 10.22, p < 0.0001). **(L-O)** During cue reinstatement, there were no differences between Veh and ABC groups in active lever presses **(L)** or cue rewards **(M)** in the No React and FR3 groups. There were also no differences between Veh and ABC groups in active lever presses for sucrose **(N)** or sucrose pellets **(O)** in the Sucrose VR5 group. **(P, Q)** Active lever presses during PR schedule over three consecutive days in No React, FR3 **(P)**, or Sucrose VR5 **(Q)** groups. A two-way RM ANOVA showed only a main effect of day for active lever presses in the Sucrose VR5 group (F_1.752,21.02_ = 10.87, p = 0.0008) and for sucrose rewards (F_1.923, 23.08_ = 12.17, p = 0.0003). **(R, S)** Microinjection sites for No React, FR3 **(R)** or Sucrose VR5 **(S)**. Gray dots = Veh, red dots = ABC. Data are mean ± SEM. No React: Veh N = 7; ABC = 6; FR3: Veh N = 5; ABC N = 6; Sucrose VR5: Veh N = 9; ABC N = 5.

**Supplemental Figure 2.**
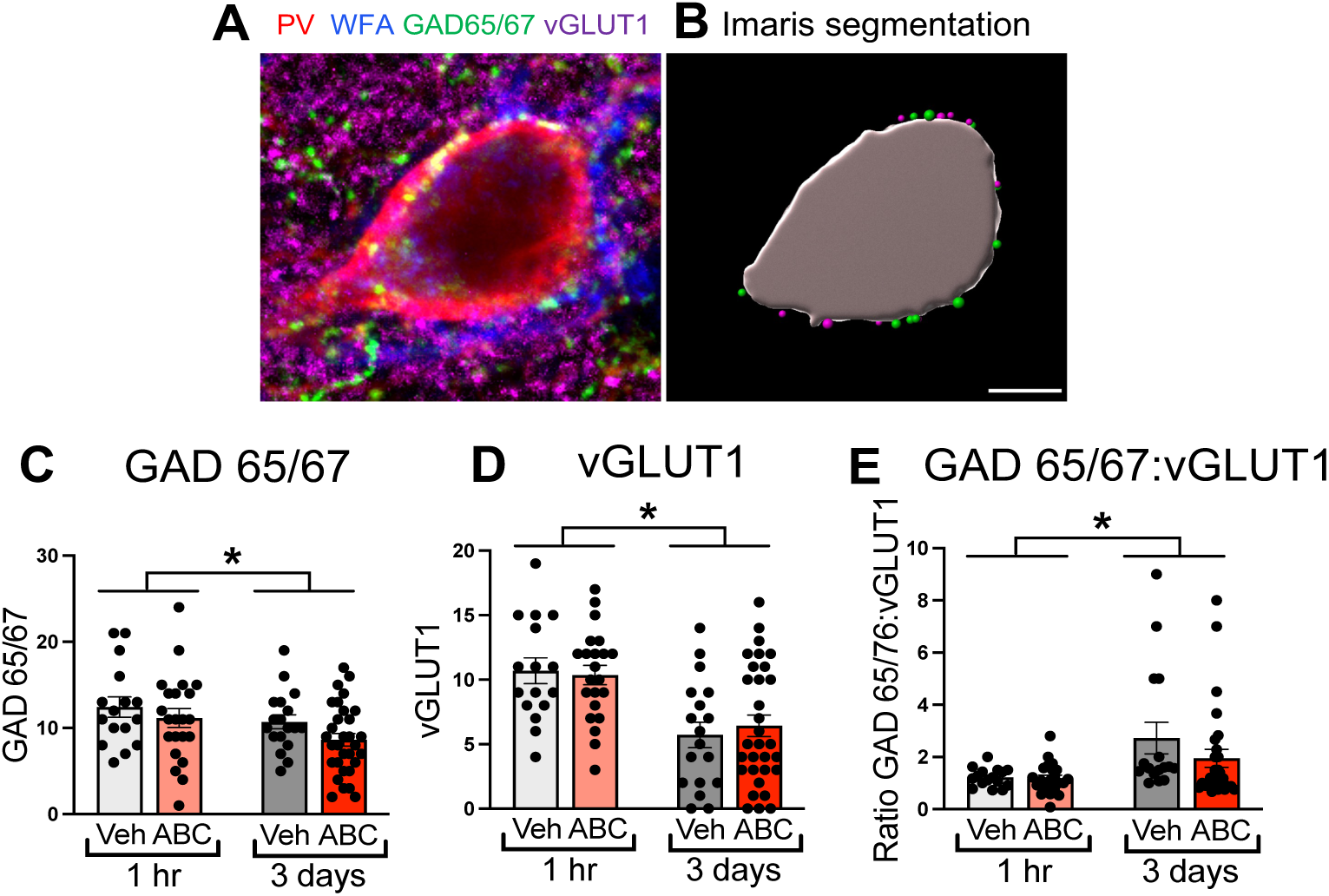
ABC treatment in the mPFC does not alter inhibitory/excitatory puncta on PV cells. **(A)** Confocal micrograph of a representative PV neuron (red) surrounded by a WFA-labeled PNN (blue). This PV neuron receives appositions from both glutamatergic (VGluT1, magenta) and GABAergic (GAD65/67, green) puncta. **(B)** The Imaris Spots segmentation tool was used to analyze GAD65/67 (green dots) and VGLUT1 (magenta dots) puncta apposing the outer boundaries of the PV neuron (gray). Scale bar = 5 µm. **(C)** GAD 65/67 puncta apposing PV neurons. There was a main effect of time (F_1, 83_ = 4.817, p = 0.031), with a reduced number of puncta at 3 days vs 1 hr. **(D)** vGLUT1 puncta apposing PV neurons. There was a main effect of time (F_1, 83_ = 23.22, p < 0.0001), with a reduced number of vGLUT1 puncta at 3 days vs. 1 hr. **(E)** Ratio of GAD65/67 to vGLUT1 puncta apposing PV neurons. There was a main effect of time (F_1, 78_ = 10.52, p = 0.0017), with a reduced ratio at 1 hr vs. 3 days. Data are mean ± SEM. Number of rats (number of cells) for each group: 1 hr: Veh N = 4 (16); ABC N = 4 (22) 3 days: Veh N = 3 (18), ABC N = 3 (31). * p < 0.05.

**Supplemental Figure 3.**
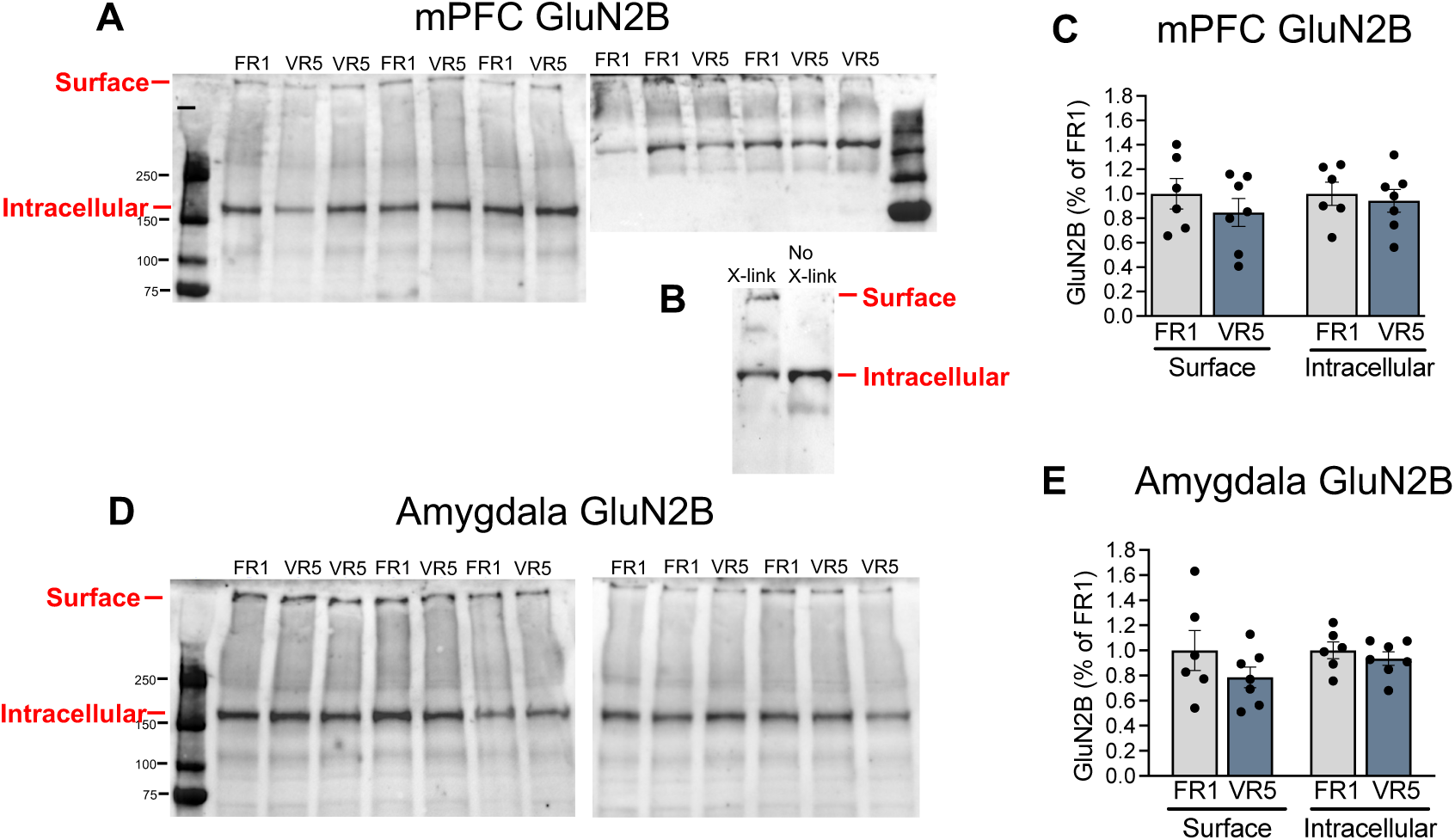
GluN2B levels are not altered in mPFC or amygdala immediately after novel VR5 memory reactivation session. **(A)** mPFC western blots showing surface and intracellular GluN2B immediately after the 30 min FR1 or VR5 memory reactivation session. **(B)** mPFC surface and intracellular mGluN2B in the absence or presence of cross-linking with BS3. **(C)** Quantification of mPFC surface and intracellular GluN2B levels after the FR1 or VR5 memory reactivation session **(D)** Amygdala western blots showing surface and intracellular GluN2B immediately after the 30 min FR1 or VR5 memory reactivation session. **(E)** Quantification of amygdala GluN2B levels after the FR1 or VR5 reactivation session. For FR1 N = 6, for VR5 N = 7.

**Supplemental Figure 4.**
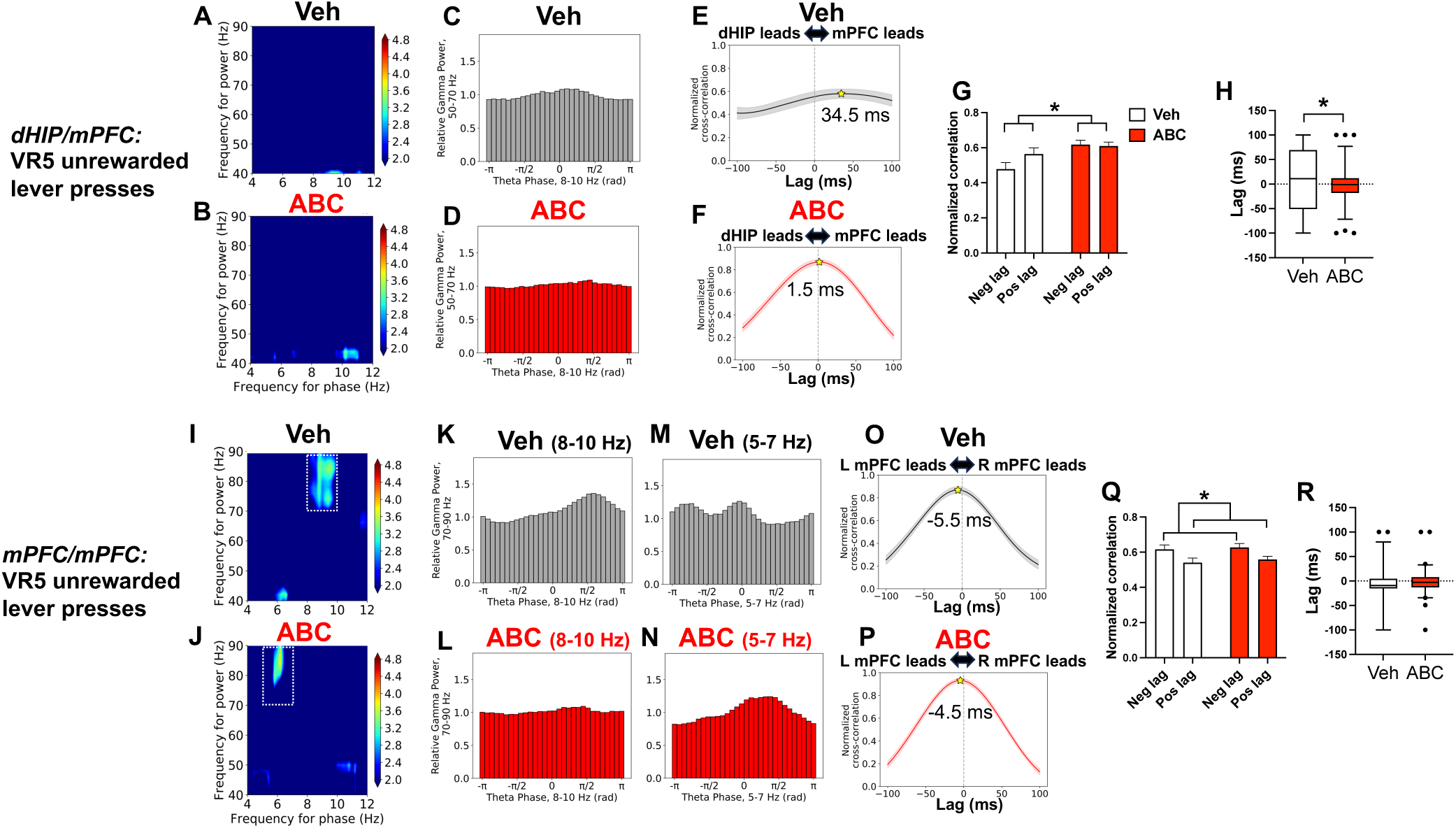
Lack of dHIP modulation of mPFC for unrewarded lever presses during novel VR5 memory reactivation, but ABC alters inter-hemispheric theta/gamma coupling across the mPFC. **(A)** Vehicle rats: comodulogram showing dHIP theta phase and mPFC amplitude 200-600 ms after unrewarded lever presses. **(B)** ABC rats: comodulogram showing dHIP theta phase and mPFC amplitude 200-600 ms after unrewarded lever presses. Color bar is z-score; comodulograms are floored at p < 0.05. **(C, D)** Vehicle **(C)** and ABC **(D)** rats: plot of mPFC gamma power (50-70 Hz) binned by dHIP theta phase (8-10 Hz) 200-600 ms after unrewarded lever presses. dHIP theta does not modulate mPFC gamma power in either Veh or ABC rats (MI = 4.89 x 10^-4^; z = 0.24 and MI = 1.50 x 10^-4^; z = −0.86 for C and D, respectively). **(E, F)** Cross-correlation between mPFC/dHIP theta (4-12 Hz) amplitude envelopes at varying lags in Veh **(E)** and ABC **(F)** groups during VR5 memory reactivation session. Stars show peak correlation (Mean ± SEM: Veh = 34.5 ± 0.005 ms; ABC = 1.5 ± 0.003 ms). Number of rats (number of trials): Veh N = 4 (59); ABC N = 3 (71). **(G)** Normalized correlation for negative (Neg) and positive (Pos) lag, with positive lag indicating mPFC leading the dHIP. There was a main effect of treatment (F_1,256_ = 9.736; p = 0.0020). **(H)** Lag at peak correlation, with a significant difference in distributions between the Veh and ABC groups (Komogorov-Smirnov p = 0.0018). **(I)** Veh rats: comodulogram showing left mPFC theta phase and right mPFC amplitude 200-600 ms after unrewarded lever presses. **(J)** ABC rats: comodulogram showing left mPFC theta phase and right mPFC amplitude 200-600 ms after unrewarded lever presses. Color bar is z-score; comodulograms are floored at p < 0.05. **(K)** Vehicle rats: plot of mPFC gamma power (70-90 Hz) binned by dHIP theta phase (8-10 Hz) 200-600 ms after unrewarded lever presses. Left mPFC theta significantly modulates right mPFC gamma power (MI = 2.38 x 10^-3^; z = 3.18). **(L)** ABC rats: plot of mPFC gamma power (70-90 Hz) binned by dHIP theta phase (8-10 Hz) 200-600 ms after unrewarded lever presses. Left mPFC theta does not modulate right mPFC gamma power after lever presses (MI = 1.67 x 10^-4^; z = −0.71). **(M)** Veh rats: plot of mPFC gamma power (70-90 Hz) binned by dHIP theta phase (5-7 Hz) 200-600 ms after unrewarded lever presses. Left mPFC theta does not modulate right mPFC gamma power (MI = 1.62 x 10^-3^; z = 0.41). **(N)** ABC rats: plot of mPFC gamma power (70-90 Hz) binned by dHIP theta phase (5-7 Hz) 200-600 ms after unrewarded lever presses. Left mPFC theta modulates right mPFC gamma power (MI = 3.04 x 10^-3^; z = 4.81). **(O, P)** Cross-correlation between left mPFC/right mPFC theta (4-12 Hz) amplitude envelopes at varying lags in Veh **(O)** and ABC **(P)** groups. Stars show peak correlation (mean ± SEM: Veh = −5.5 ms ± 0.004 ms; ABC = −4.5 ± 0.003 ms). Line is mean, shading represents SEM. **(Q)** Normalized correlation for negative (Neg) and positive (Pos) lag, with positive lag indicating left mPFC leading the right mPFC. There was a main effect of lag (F_1,232_) = 10.57; p = 0.0013). **(R)** Lag at peak correlation, with no difference in distributions between the Veh and ABC groups (Komogorov-Smirnov p = 0.4025). Box plot shows median and 95% confidence intervals.

## Notes

### Competing Interest Statement

The authors have declared no competing interest.

