## Supplemental Table 1 for "Perineuronal nets in the rat medial prefrontal cortex alter hippocampal-prefrontal oscillations and reshape cocaine self-administration memories"

**Supplemental Table 1. Cocaine or sucrose self-administration total active and inactive lever presses and total rewards**

| TREATMENT GROUP (n) | Total active lever presses | Total rewards | Total inactive lever presses |
|---|---|---|---|
| Acquisition Veh (8) | 1952 ± 182 | 1431 ± 153 | 353 ± 99 |
| Acquisition ABC(6) | **966 ± 242*** | **702 ± 190*** | **208 ± 89*** |
| FR1 Veh (9) | 343 ± 34 | 234 ± 20 | 122 ± 53 |
| FR1 ABC (6) | 535 ± 93 | 312 ± 24 | 94 ± 54 |
| Pre-VR5 Veh (15) | 525 ± 102 | 265 ± 25 | 59 ± 10 |
| Pre-VR5 ABC (10) | 411 ± 73 | 232 ± 28 | 63 ± 28 |
| Post-VR5 Veh (9) | 750 ± 261 | 282 ± 23 | 133 ± 41 |
| Post-VR5 ABC (7) | 621 ± 150 | 292 ± 43 | 919 ± 839 |
| No React Veh (7) | 447 ± 104 | 301 ± 54 | 126 ± 66 |
| No React ABC (6) | 631 ± 222 | 344 ± 82 | 1601 ± 1322 |
| FR3 Veh (5) | 726 ± 196 | 242 ± 40 | 62 ± 15 |
| FR3 ABC (6) | 981 ± 307 | 272 ± 43 | 511 ± 473 |
| Sucrose Veh (9) | 821 ± 122 | 603 ± 80 | 128 ± 26 |
| Sucrose ABC (5) | 670 ± 118 | 482 ± 74 | 126 ± 21 |

Data are mean ± SEM. Total rewards refer to both cocaine infusions and sucrose pellets consumed. *$P < 0.05$, compared with Veh group using a two-way RM ANOVA.
