## Supplemental Table 2 for "Perineuronal nets in the rat medial prefrontal cortex alter hippocampal-prefrontal oscillations and reshape cocaine self-administration memories"

### Supplemental Table 2. Statistical comparisons: Figure 1

| Figure | Measure | Groups | N-size | Test | F | p value |
|---|---|---|---|---|---|---|
| 1B | Acquisition: Active lever | Veh | 8 | 2-way RM ANOVA | Treatment F (1, 12) = 11.10 | **0.0006** |
| | | ABC | 6 | | Day F (3.22, 38.58) = 8.815 | **< 0.0001** |
| | | | | | Interaction F (19, 228) = 1.147 | 0.3055 |
| 1C | Acquisition: Infusions | Veh | 8 | 2-way RM ANOVA | Treatment F (1, 12) = 9.167 | **0.0105** |
| | | ABC | 6 | | Day F (2.9, 31.14) = 10.71 | **< 0.0001** |
| | | | | | Interaction F (19, 228) = 0.8813 | 0.6066 |
| 1D | Acquisition: Inactive lever | Veh | 8 | 2-way RM ANOVA | Treatment F (1, 12) = 1.122 | 0.3104 |
| | | ABC | 6 | | Day F (2.44, 29.23) = 0.8382 | 0.4629 |
| | | | | | Interaction F (19, 228) = 0.3673 | 0.9935 |
