## Supplemental Table 3 for "Perineuronal nets in the rat medial prefrontal cortex alter hippocampal-prefrontal oscillations and reshape cocaine self-administration memories"

**Supplemental Table 3. Figure 2 rewards and inactive lever presses**

| Associated Figure | Measure | Groups | N-size | Day 8 mean ± SEM | Day 9 mean ± SEM | Day 10 mean ± SEM |
|---|---|---|---|---|---|---|
| 2B,C | Last 3 days training: Inactive lever presses | FR1 Veh | 9 | 4 ± 2 | 3 ± 1 | 55 ± 54 |
| | | FR1 ABC | 6 | 2 ± 1 | 1 ± 1 | 1 ± 1 |
| | | Pre-VR5 Veh | 15 | 4 ± 2 | 3 ± 1 | 3 ± 1 |
| | | Pre-VR5 ABC | 10 | 4 ± 3 | 2 ± 1 | 3 ± 1 |
| | | Post-VR5 Veh | 9 | 8 ± 5 | 3 ± 2 | 2 ± 1 |
| | | Post-VR5 ABC | 7 | 7 ± 4 | 1 ± 1 | 1 ± 1 |
| 2D,E | Memory Reactivate: Inactive lever presses | FR1 Veh | 9 | 9 ± 8 | | |
| | | FR1 ABC | 6 | 1 ± 1 | | |
| | | Pre-VR5 Veh | 15 | 3 ± 1 | | |
| | | Pre-VR5 ABC | 10 | 3 ± 1 | | |
| | | Post-VR5 Veh | 9 | 2 ± 1 | | |
| | | Post-VR5 ABC | 7 | 1 ± 1 | | |
| 2F | Extinction FR1: Inactive lever presses | **Time course** | | **5 10 15 20 25 30 min** | | |
| | | Veh | 9 | 1±1, 1±1, 1±1, 2±1, 1±1, 1±1 | | |
| | | ABC | 6 | 2±1, 2±1, 2±1, 1±1, 1±1, 1±1 | | |
| 2G | Extinction Pre-VR5: Inactive lever presses | **Time course** | | **5 10 15 20 25 30 min** | | |
| | | Veh | 15 | 2±1, 1±1, 1±1, 1±1, 1±1, 1±1 | | |
| | | ABC | 10 | 1±1, 1±1, 1±1, 0±0, 0±0, 0±0 | | |
| 2H | Extinction Post-VR5: Inactive lever presses | **Time course** | | **5 10 15 20 25 30 min** | | |
| | | Veh | 9 | 2±1, 2±1, 2±1, 1±1, 1±1, 2±1 | | |
| | | ABC | 7 | 7±5, 2±1, 2±1, 2±1, 1±1, 1±1 | | |
| 2I,J | Cue Reinstatement: Inactive lever presses | FR1 Veh | 9 | 3 ± 1 | | |
| | | FR1 ABC | 6 | 5 ± 3 | | |
| | | Pre-VR5 Veh | 15 | 3 ± 1 | | |
| | | Pre-VR5 ABC | 10 | 1 ± 1 | | |
| | | Post-VR5 Veh | 9 | 4 ± 1 | | |
| | | Post-VR5 ABC | 7 | 6 ± 2 | | |
| 2K | Progressive Ratio: Rewards | | | **Day 1** | **Day 2** | **Day 3** |
| | | FR1 Veh | 9 | 13 ± 1 | 14 ± 1 | 14 ± 1 |
| | | FR1 ABC | 6 | 12 ± 2 | 12 ± 2 | 13 ± 2 |
| | | Pre-VR5 Veh | 10 | 11 ± 1 | 11 ± 2 | 11 ± 1 |
| | | Pre-VR5 ABC | 7 | 7 ± 1 | 7 ± 2 | 7 ± 2 |
| 2K | Progressive Ratio: Inactive lever presses | | | **Day 1** | **Day 2** | **Day 3** |
| | | FR1 Veh | 9 | 23 ± 5 | 21 ± 6 | 21 ± 7 |
| | | FR1 ABC | 6 | 31 ± 13 | 23 ± 9 | 18 ± 8 |
| | | Pre-VR5 Veh | 10 | 17 ± 5 | 16 ± 4 | 13 ± 3 |
| | | Pre-VR5 ABC | 7 | 4 ± 1 | 3 ± 1 | 2 ± 1 |
