## Supplemental Table 4 for "Perineuronal nets in the rat medial prefrontal cortex alter hippocampal-prefrontal oscillations and reshape cocaine self-administration memories"

**Supplemental Table 4. Statistical comparisons: Figure 2**

| Figure | Measure | Groups | N-size | Test | F | p value | Šídák's | p value |
|---|---|---|---|---|---|---|---|---|
| 2B | Last 3 days training: Active lever presses | FR1 Veh | 9 | 3-way RM ANOVA | Treatment (Veh vs ABC) F (1,50) = 0.182 | 0.671 | | |
| | | FR1 ABC | 6 | | React F (2, 50) = 1.389 | 0.259 | | |
| | | Pre-VR5 Veh | 15 | | Day F (1.28, 64.18) = 1.414 | 0.246 | | |
| | | Pre-VR5 ABC | 10 | | Treatment x React F (2, 49) = 0.331 | 0.720 | | |
| | | Post-VR5 Veh | 9 | | Treatment x Day F (1.28, 64.18) = 1.199 | 0.291 | | |
| | | Post-VR5 ABC | 7 | | React x Day F (2.57, 64.18) = 0.329 | 0.773 | | |
| | | | | | Treatment x React x Day F (2.57, 64.18) = 1.464 | 0.236 | | |
| 2C | Last 3 days training: Cocaine infusions | FR1 Veh | 9 | 3-way RM ANOVA | Treatment (Veh vs ABC) F (1, 50) = 1.027 | 0.316 | | |
| | | FR1 ABC | 6 | | React F (2, 50) = 0.388 | 0.680 | | |
| | | Pre-VR5 Veh | 15 | | Day F (1.73, 86.35) = 0.055 | 0.926 | | |
| | | Pre-VR5 ABC | 10 | | Treatment x React F (2, 50) = 1.710 | 0.191 | | |
| | | Post-VR5 Veh | 9 | | Treatment x Day F (1.73, 86.35) = 0.181 | 0.803 | | |
| | | Post-VR5 ABC | 7 | | React x Day F (3.45, 86.35) = 1.070 | 0.371 | | |
| | | | | | Treatment x React x Day F (3.45, 86.35) = 0.975 | 0.417 | | |
| Suppl T3 | Last 3 days training: Inactive lever presses | FR1 Veh | 9 | 3-way RM ANOVA | Treatment (Veh vs ABC) F (1, 50) = 1.390 | 0.244 | | |
| | | FR1 ABC | 6 | | React F (2, 50) = 0.672 | 0.515 | | |
| | | Pre-VR5 Veh | 15 | | Day F (1.03, 51.38) = 0.720 | 0.404 | | |
| | | Pre-VR5 ABC | 10 | | Treatment x React F (2, 49) = 0.981 | 0.382 | | |
| | | Post-VR5 Veh | 9 | | Treatment x Day F (1.03, 51.38) = 0.893 | 0.352 | | |
| | | Post-VR5 ABC | 7 | | React x Day F (2.05, 51.38) = 0.936 | 0.401 | | |
| | | | | | Treatment x React x Day F (2.05, 51.38) = 0.863 | 0.431 | | |
| 2D | Memory Reactivate: Active lever presses | FR1 Veh/ABC | 9/6 | 2-way ANOVA | Treatment (Veh vs ABC) F (1, 50) = 0.2485 | 0.6203 | | |
| | | Pre-VR5 Veh/ABC | 15/10 | | React (FR1 vs VR5 (pre/post) F (2, 50) = 23.98 | **< 0.0001** | FR1 vs Pre-VR5 | **< 0.0001** |
| | | Post-VR5 Veh/ABC | 9/7 | | Treatment x React F (2, 50) = 0.2063 | 0.8142 | FR1 vs. Post-VR5 | **< 0.0001** |

**Supplemental Table 4. Statistical comparisons: Figure 2 - con't**

| Figure | Measure | Groups | N-size | Test | F | p value | Šídák's | p value |
|---|---|---|---|---|---|---|---|---|
| 2E | Memory Reactivate: Cocaine infusions | FR1 Veh/ABC | 9/6 | 2-way ANOVA | Treatment (Veh vs ABC) F (1,50) = 0.06458 | 0.8004 | | |
| | | Pre-VR5 Veh/ABC | 15/10 | | React (FR1 vs VR5 (pre/post) F (2, 50) = 8.136 | **0.0009** | FR1 vs Pre-VR5 | **0.0003** |
| | | Post-VR5 Veh/ABC | 9/7 | | Treatment x React F (2, 50) = 0.2349 | 0.7915 | | |
| Suppl T3 | Memory Reactivate: Inactive lever presses | FR1 Veh/ABC | 9/6 | 2-way ANOVA | Treatment (Veh vs ABC) F (1, 50) = 0.9371 | 0.3377 | | |
| | | Pre-VR5 Veh/ABC | 15/10 | | React (FR1 vs VR5 (pre/post) F (2, 50) = 0.3753 | 0.6890 | | |
| | | Post-VR5 Veh/ABC | 9/7 | | Treatment x React F (2, 50) = 0.8527 | 0.4324 | | |
| 2F | Extinction FR1: Active lever presses | Veh | 9 | 2-way RM ANOVA | Treatment F (1, 13) = 0.3662 | 0.8512 | | |
| | | ABC | 6 | | Time F (3.196, 41.54) = 7.467 | **0.0003** | | |
| | | | | | Treatment x Time F (5, 65) = 0.3878 | 0.8554 | | |
| Suppl T3 | Extinction FR1: Inactive lever presses | Veh | 9 | 2-way RM ANOVA | Treatment F (1, 13) = 0.7220 | 0.4109 | | |
| | | ABC | 6 | | Time F (3.151, 40.96) = 1.649 | 0.1912 | | |
| | | | | | Treatment x Time F (5, 65) = 0.7017 | 0.6241 | | |
| 2G | Extinction Pre-VR5: Active lever presses | Veh | 15 | 2-way RM ANOVA | Treatment F (1, 23) = 2.608 | 0.1200 | | |
| | | ABC | 10 | | Time F (3.583, 82.40) = 9.779 | **< 0.0001** | | |
| | | | | | Treatment x Time F (5, 115) = 0.5818 | 0.7139 | | |
| Suppl T3 | Extinction Pre-VR5: Inactive lever presses | Veh | 15 | 2-way RM ANOVA | Treatment F (1, 23) = 5.394 | **0.0294** | | |
| | | ABC | 10 | | Time F (2.042, 46.97) = 2.880 | 0.0650 | | |
| | | | | | Treatment x Time F (5, 115) = 1.492 | 0.1979 | | |

**Supplemental Table 4. Statistical comparisons: Figure 2 - con't**

| Figure | Measure | Groups | N-size | Test | F | p value | Šídák's | p value |
|---|---|---|---|---|---|---|---|---|
| 2H | Extinction Post-VR5: Active lever presses | Veh | 9 | 2-way RM ANOVA | Treatment F (1, 7) = 1.608 | 0.2254 | | |
| | | ABC | 7 | | Time F (3.081, 43.13) = 13.79 | **< 0.0001** | | |
| | | | | | Treatment x Time F (5, 70) = 1.899 | 0.1055 | | |
| Suppl T3 | Extinction Post-VR5: Inactive lever presses | Veh | 9 | 2-way RM ANOVA | Treatment F (1, 14) = 0.8548 | 0.3709 | | |
| | | ABC | 7 | | Time F (1.723, 24.12) = 1.540 | 0.2347 | | |
| | | | | | Treatment x Time F (5, 70) = 1.047 | 0.3973 | | |
| 2I | Cue Reinstatement: Active lever presses | FR1 Veh/ABC | 9/6 | 2-way ANOVA | Treatment (Veh vs ABC) F (1, 50) = 0.4069 | 0.5265 | | |
| | | Pre-VR5 Veh/ABC | 15/10 | | React FR1 vs VR5 (pre/post) F (2, 50) = 9.303 | **0.0004** | Pre-VR5:Veh vs. ABC | **0.0079** |
| | | Post-VR5 Veh/ABC | 9/7 | | Treatment x React F (2, 50) = 6.145 | **0.0041** | React ABC:Pre- vs. Post-VR5 | **< 0.0001** |
| 2J | Cue Reinstatement: Cue rewards | FR1 Veh/ABC | 9/6 | 2-way ANOVA | Treatment (Veh vs ABC) F (1, 50) = 0.5452 | 0.4637 | | |
| | | Pre-VR5 Veh/ABC | 15/10 | | React (FR1 vs VR5 (pre/post) F (2, 50) = 8.612 | **0.0006** | Pre-VR5: Veh vs. ABC | **0.0233** |
| | | Post-VR5 Veh/ABC | 9/7 | | Treatment x React F (2, 50) = 4.266 | **0.0195** | React ABC:FR1 vs. Pre-VR5 | **0.0167** |
| | | | | | | | React ABC:Pre- vs. Post VR5 | **0.0002** |
| Suppl T3 | Cue Reinstatement: Inactive lever presses | FR1 Veh/ABC | 9/6 | 2-way ANOVA | Treatment (Veh vs ABC) F (1, 50) = 0.2352 | 0.6298 | | |
| | | Pre-VR5 Veh/ABC | 15/10 | | React (FR1 vs VR5 (pre/post) F (2, 50) = 4.461 | **0.0165** | | |
| | | Post-VR5 Veh/ABC | 9/7 | | Treatment x React F (2, 50) = 1.686 | 0.1957 | | |
| 2K | FR1 vs VR5 PR: Active lever presses | FR1 Veh | 9 | 3-way RM ANOVA | Treatment (Veh vs ABC) F (1, 28) = 0.945 | 0.339 | | |
| | | FR1 ABC | 6 | | React F (1, 28) = 4.665 | **0.039** | | |
| | | Pre-VR5 Veh | 10 | | Day F (1.84, 51.41) = 2.089 | 0.138 | | |
| | | Pre-VR5 ABC | 7 | | Treatment x React F (1, 28) = 0.288 | 0.596 | | |
| | | | | | Treatment x Day F (1.84, 51.41) = 0.600 | 0.539 | | |
| | | | | | React x Day F (1.84, 51.41) = 1.024 | 0.361 | | |
| | | | | | Treatment x React x Day F (1.84, 51.41) = 0.220 | 0.785 | | |

### Supplemental Table 4. Statistical comparisons: Figure 2 - con't

| Figure | Measure | Groups | N-size | Test | F | p value | Šídák's | p value |
|---|---|---|---|---|---|---|---|---|
| Suppl T3 | FR1 vs VR5 PR: Cocaine infusions | FR1 Veh | 9 | 3-way RM ANOVA | Treatment F (1, 28) = 3.099 | 0.089 | | |
| | | FR1 ABC | 6 | | React F (1, 28) = 7.438 | **0.011** | | |
| | | Pre-VR5 Veh | 10 | | Day F (1.8, 50.28) = 2.049 | 0.144 | | |
| | | Pre-VR5 ABC | 7 | | Treatment x React F (1, 28) = 0.601 | 0.445 | | |
| | | | | | Treatment x Day F (1.8, 50.28) = 0.655 | 0.508 | | |
| | | | | | React x Day F (1.8, 50.28) = 0.997 | 0.369 | | |
| | | | | | Treatment x React x Day F (1.8, 50.28) = 0.335 | 0.694 | | |
| Suppl T3 | FR1 vs VR5 PR: Inactive lever presses | FR1 Veh | 9 | 3-way RM ANOVA | Treatment F (1, 28) = 1.126 | 0.298 | | |
| | | FR1 ABC | 6 | | React F (1, 28) = 8.844 | **0.006** | | |
| | | Pre-VR5 Veh | 10 | | Day F (1.53, 42.8) = 1.696 | 0.200 | | |
| | | Pre-VR5 ABC | 7 | | Treatment x React F (1, 28) = 2.431 | 0.130 | | |
| | | | | | Treatment x Day F (1.53, 42.8) = 0.282 | 0.696 | | |
| | | | | | React x Day F (1.53, 42.8) = 0.341 | 0.655 | | |
| | | | | | Treatment x React x Day F (1.53, 42.8) = 0.791 | 0.429 | | |
