## Supplemental Table 5 for "Perineuronal nets in the rat medial prefrontal cortex alter hippocampal-prefrontal oscillations and reshape cocaine self-administration memories"

### Supplemental Table 5. Supplemental Figure 1 rewards and inactive lever presses

| Associated Figure | Measure | Groups | N-size | Day 8 mean ± SEM | Day 9 mean ± SEM | Day 10 mean ± SEM |
|---|---|---|---|---|---|---|
| Suppl 1A-D | Last 3 days training: Inactive lever presses | NR Veh | 7 | 3 ± 2 | 2 ± 1 | 4 ± 2 |
| | | NR ABC | 6 | 292 ± 291 | 270 ± 269 | 169± 169 |
| | | FR3 Veh | 5 | 1 ± 1 | 4 ± 3 | 3 ± 3 |
| | | FR3 ABC | 6 | 1 ± 1 | 1 ± 1 | 1 ± 1 |
| | | Sucrose Veh | 9 | 12 ± 4 | 11 ± 3 | 13 ± 5 |
| | | Sucrose ABC | 5 | 15 ± 6 | 12 ± 4 | 16 ± 3 |
| Suppl 1E-H | Memory Reactivate: Inactive lever presses | FR3 Veh | 5 | 1 ± 1 | | |
| | | FR3 ABC | 6 | 1 ± 1 | | |
| | | Sucrose Veh | 9 | 13 ± 3 | | |
| | | Sucrose ABC | 5 | 7 ± 3 | | |
| Suppl 1I | Extinction NR: Inactive lever presses | **Time course** | | **5 10 15 20 25 30 min** | | |
| | | Veh | 7 | 1±1, 2±1, 1±1, 1±1, 1±1, 1±1 | | |
| | | ABC | 6 | 3±2, 4±1, 3±1, 1±1, 1±1, 0±0 | | |
| Suppl 1J | Extinction FR3: Inactive lever presses | **Time course** | | **5 10 15 20 25 30 min** | | |
| | | Veh | 5 | 2±1, 2±1, 2±1, 2±1, 0±1, 0±1 | | |
| | | ABC | 6 | 2±1, 1±1, 2±1, 0±1, 1±1, 1±1 | | |
| Suppl 1K | Extinction Sucrose: Inactive lever presses | **Time course** | | **5 10 15 20 25 30 min** | | |
| | | Veh | 9 | 2±1, 3±1, 1±1, 1±1, 1±1, 1±1 | | |
| | | ABC | 5 | 3±1, 1±1, 2±1, 1±1, 1±1, 1±1 | | |
| Suppl 1L-O | Cue Reinstatement: Inactive lever presses | NR Veh | 7 | 3 ± 3 | | |
| | | NR ABC | 6 | 3 ± 2 | | |
| | | FR3 Veh | 5 | 4 ± 1 | | |
| | | FR3 ABC | 6 | 2 ± 1 | | |
| | | Sucrose Veh | 9 | 3 ± 2 | | |
| | | Sucrose ABC | 5 | 6 ± 2 | | |
| Suppl 1 P,Q | Progressive Ratio: Rewards | | | **Day 1** | **Day 2** | **Day 3** |
| | | NR Veh | 7 | 13 ± 1 | 14 ± 1 | 14 ± 1 |
| | | NR ABC | 6 | 12 ± 2 | 12 ± 2 | 12 ± 2 |
| | | FR3 Veh | 5 | 12 ± 1 | 12 ± 2 | 13 ± 2 |
| | | FR3 ABC | 6 | 13 ± 1 | 15 ± 1 | 14 ± 1 |
| | | Sucrose Veh | 9 | 10 ± 1 | 9 ± 1 | 8 ± 1 |
| | | Sucrose ABC | 5 | 11 ± 1 | 10 ± 1 | 9 ± 1 |
| Suppl 1 P,Q | Progressive Ratio: Inactive lever presses | | | **Day 1** | **Day 2** | **Day 3** |
| | | NR Veh | 7 | 27 ± 11 | 20 ± 6 | 14 ± 6 |
| | | NR ABC | 6 | 17 ± 7 | 14 ± 5 | 19 ± 7 |
| | | FR3 Veh | 5 | 23 ± 6 | 27 ± 7 | 30 ± 5 |
| | | FR3 ABC | 6 | 10 ± 2 | 17 ± 4 | 18 ± 5 |
| | | Sucrose Veh | 9 | 21 ± 8 | 24 ± 12 | 20 ± 10 |
| | | Sucrose ABC | 5 | 25 ± 8 | 16 ± 4 | 17 ± 5 |
