## Supplemental Table 6 for "Perineuronal nets in the rat medial prefrontal cortex alter hippocampal-prefrontal oscillations and reshape cocaine self-administration memories"

**Supplemental Table 6: Statistical comparisons: Supplemental Figure 1**

| Figure | Measure | Groups | N-size | Test | F | p value |
|---|---|---|---|---|---|---|
| Suppl 1A | Last 3 days training: Active lever presses | NR Veh | 7 | 3-way RM ANOVA | Treatment (Veh vs ABC) F (1,20) = 0.130 | 0.722 |
| | | NR ABC | 6 | | React F (1, 20) = 9.426 | **0.006** |
| | | FR3 Veh | 5 | | Day F (1.31, 26.20) = 1.782 | 0.194 |
| | | FR3 ABC | 6 | | Treatment x React F (1, 20) = 0.278 | 0.604 |
| | | | | | Treatment x Day F (1.31, 26.20) = 0.137 | 0.782 |
| | | | | | React x Day F (1.31, 26.20) = 0.461 | 0.555 |
| | | | | | Treatment x React x Day F (1.31, 26.20)=2.128 | 0.152 |
| Suppl 1B | Last 3 days training: Cocaine infusions | NR Veh | 7 | 3-way RM ANOVA | Treatment (Veh vs ABC) F (1, 20) = 0.122 | 0.731 |
| | | NR ABC | 6 | | React F (1, 20) = 2.441 | 0.134 |
| | | FR3 Veh | 5 | | Day F (1.23, 24.63) = 1.779 | 0.195 |
| | | FR3 ABC | 6 | | Treatment x React F (1, 20) = 0.085 | 0.774 |
| | | | | | Treatment x Day F (1.23, 24.63) = 0.316 | 0.625 |
| | | | | | React x Day F (1.23, 24.63) = 1.312 | 0.272 |
| | | | | | Treatment x React x Day F (1.23, 24.63)=2.863 | 0.097 |
| Suppl T5 | Last 3 days training: Inactive lever presses | NR Veh | 7 | 3-way RM ANOVA | Treatment (Veh vs ABC) F (1, 20) = 0.954 | 0.34 |
| | | NR ABC | 6 | | React F (2, 20) = 0.982 | 0.334 |
| | | FR3 Veh | 5 | | Day F (1, 20.01) = 0.918 | 0.35 |
| | | FR3 ABC | 6 | | Treatment x React F (1, 20) = 0.978 | 0.334 |
| | | | | | Treatment x Day F (1, 20.01) = 1.046 | 0.319 |
| | | | | | React x Day F (1, 20.01) = 0.984 | 0.333 |
| | | | | | Treatment x React x Day F (1, 20.01)=1.002 | 0.329 |
| Suppl 1C | Last 3 days training: Sucrose Active lever presses | Veh | 9 | 2-way RM ANOVA | Treatment (Veh vs ABC) F (1, 12) = 0.3269 | 0.5780 |
| | | ABC | 5 | | Day F (1.961, 23.54) = .6350 | 0.5358 |
| | | | | | Treatment x Day F (2, 24) = 0.002225 | 0.9978 |
| Suppl 1D | Last 3 days training: Sucrose pellets | Veh | 9 | 2-way RM ANOVA | Treatment (Veh vs ABC) F (1, 12) = 0.2491 | 0.6267 |
| | | ABC | 5 | | Day F (1.890, 22.68) = 1.064 | 0.3582 |
| | | | | | Treatment x Day F (2, 24) = 0.2333 | 0.7937 |

**Supplemental Table 6: Statistical comparisons: Supplemental Figure 1-con't**

| Figure | Measure | Groups | N-size | Test | F | p value |
|---|---|---|---|---|---|---|
| Suppl T5 | Last 3 days training: Sucrose Inctive lever presses | Veh/ABC | 9 | 2-way RM ANOVA | Treatment (Veh vs ABC) F (1, 12) = 0.1540 | 0.7016 |
| | | Veh/ABC | 5 | | Day F (1.876, 22.51) = 0.7758 | 0.4646 |
| | | | | | Treatment x Day F (2, 24) = 0.2062 | 0.8151 |
| Suppl 1E | Memory Reactivate: FR3 Active lever presses | Veh | 5 | Student's t-test | | |
| | | ABC | 6 | | t = 0.0413, df = 9 | 0.9679 |
| Suppl 1F | Memory Reactivate: FR3 Cocaine infusions | Veh | 5 | Student's t-test | | |
| | | ABC | 6 | | t = 0.6171, df = 9 | 0.5525 |
| Suppl T5 | Memory Reactivate: FR3 Inactive lever presses | Veh | 5 | Student's t-test | | |
| | | ABC | 6 | | t = 0.8435, df = 9 | 0.4208 |
| Suppl 1G | Memory Reactivate: Sucrose Active lever | Veh | 9 | Student's t-test | | |
| | | ABC | 5 | | t = 1.083, df = 12 | 0.3002 |
| Suppl 1H | Memory Reactivate: Sucrose pellets | Veh | 9 | Student's t-test | | |
| | | ABC | 5 | | t = 1.756, df = 12 | 0.1045 |
| Suppl T5 | Memory Reactivate: Sucrose Inactive lever | Veh | 9 | Student's t-test | | |
| | | ABC | 5 | | t = 1.172, df = 12 | 0.2640 |
| Suppl 1I | Extinction NR: Active lever presses | Veh | 7 | 2-way RM ANOVA | Treatment (Veh vs ABC) F (1, 11) = 0.7199 | 0.4142 |
| | | ABC | 6 | | Time F (2.426, 26.69) = 14.01 | **< 0.0001** |
| | | | | | Treatment x Time F (5, 55) = 0.2689 | 0.9282 |

**Supplemental Table 6: Statistical comparisons: Supplemental Figure 1-con't**

| Figure | Measure | Groups | N-size | Test | F | p value |
|---|---|---|---|---|---|---|
| Suppl T5 | Extinction NR: Inactive lever presses | Veh | 7 | 2-way RM ANOVA | Treatment (Veh vs ABC) F (1, 11) = 3.276 | 0.0977 |
| | | ABC | 6 | | Time F (2.862, 31.49) = 4.316 | **0.0127** |
| | | | | | Treatment x Time F (5, 55) = 1.282 | 0.2849 |
| Suppl 1J | Extinction FR3: Active lever presses | Veh | 5 | 2-way RM ANOVA | Treatment (Veh vs ABC) F (1, 9) = 0.06215 | 0.8087 |
| | | ABC | 6 | | Time F (2.085, 18.77) = 21.25 | **< 0.0001** |
| | | | | | Treatment x Time F (5, 45) = 0.2833 | 0.9199 |
| Suppl T5 | Extinction FR3: Inactive lever presses | Veh | 5 | 2-way RM ANOVA | Treatment F (1, 9) = 2.439 | 0.1528 |
| | | ABC | 6 | | Time F (2.401, 21.61) = 1.374 | 0.2766 |
| | | | | | Interaction F (5, 45) = 0.6124 | 0.6909 |
| Suppl 1K | Extinction Sucrose: Active lever presses | Veh | 9 | 2-way RM ANOVA | Treatment (Veh vs ABC) F (1, 12) = 3.318 | 0.0935 |
| | | ABC | 5 | | Time F (3.630, 43.56) = 10.22 | **< 0.0001** |
| | | | | | Treatment x Time F (5, 60) = 1.252 | 0.2966 |
| Suppl T5 | Extinction Sucrose: Inactive lever presses | Veh | 9 | 2-way RM ANOVA | Treatment (Veh vs ABC) F (1, 12) = 0.01165 | 0.9158 |
| | | ABC | 5 | | Time F (1.960, 23.52) = 3.364 | 0.0527 |
| | | | | | Treatment x Time F (5, 60) = 0.9359 | 0.4644 |
| Suppl 1L | Cue Reinstatement: Active lever presses | NR Veh | 7 | 2-way ANOVA | Treatment (Veh vs ABC) F (1, 20) = 0.2714 | 0.6081 |
| | | NR ABC | 6 | | Time F (1, 20) = 0.0724 | 0.7906 |
| | | FR3 Veh | 5 | | Treatment x Time F (1, 20) = 0.00177 | 0.9668 |
| | | FR3 ABC | 6 | | | |
| Suppl 1M | Cue Reinstatement: Cue rewards | NR Veh | 7 | 2-way ANOVA | Treatment (Veh vs ABC) F (1, 20) = 0.6553 | 0.4277 |
| | | NR ABC | 6 | | Time F (1, 20) = 0.2901 | 0.5961 |
| | | FR3 Veh | 5 | | Treatment x Time F (1, 20) = 0.1958 | 0.6629 |
| | | FR3 ABC | 6 | | | |

**Supplemental Table 6: Statistical comparisons: Supplemental Figure 1-con't**

| Figure | Measure | Groups | N-size | Test | F | p value |
|---|---|---|---|---|---|---|
| Suppl T5 | Cue Reinstatement: Inactive lever presses | NR Veh | 7 | 2-way ANOVA | Treatment (Veh vs ABC) F (1, 20) = 0.2426 | 0.6277 |
| | | NR ABC | 6 | | Time F (1, 20) = 0.003294 | 0.9548 |
| | | FR3 Veh | 5 | | Treatment x Time F (1, 20) = 0.1978 | 0.6613 |
| | | FR3 ABC | 6 | | | |
| Suppl 1N | Cue Reinstatement: Active lever presses | Sucrose Veh | 9 | Student's t-test | | |
| | | Sucrose ABC | 5 | | t = 0.9499, df = 12 | 0.3609 |
| Suppl 1O | Cue Reinstatement: Cue rewards | Sucrose Veh | 9 | Student's t-test | | |
| | | Sucrose ABC | 5 | | t = 1.005, df = 12 | 0.3346 |
| Suppl T5 | Cue Reinstatement: Inactive lever presses | Sucrose Veh | 9 | Student's t-test | | |
| | | Sucrose ABC | 5 | | t = 0.9430, df = 12 | 0.3643 |
| Suppl 1P | NR vs FR3 PR: Active lever presses | NR Veh | 7 | 3-way RM ANOVA | Treatment F (1, 20) = 0.0880 | 0.769 |
| | | NR ABC | 6 | | React F (1, 20) = 1.57 e-5 | 0.997 |
| | | FR3 Veh | 5 | | Day F (1.68, 33.53) = 3.386 | 0.054 |
| | | FR3 ABC | 6 | | Treatment x React F (1, 20) = 0.7389 | 0.400 |
| | | | | | Treatment x Day F (1.68, 33.53) = 1.580 | 0.222 |
| | | | | | React x Day F (1.68, 33.53) = 1.197 | 0.308 |
| | | | | | Treatment x React x Day F (1.68, 33.53)=0.424 | 0.622 |
| Suppl T5 | NR vs FR3 PR: Cocaine infusions | NR Veh | 7 | 3-way RM ANOVA | Treatment F (1, 20) = 0.0360 | 0.851 |
| | | NR ABC | 6 | | React F (1, 20) = 0.354 | 0.558 |
| | | FR3 Veh | 5 | | Day F (1.86, 37.11) = 2.334 | 0.114 |
| | | FR3 ABC | 6 | | Treatment x React F (1, 20) = 1.425 | 0.247 |
| | | | | | Treatment x Day F (1.86, 37.11) = 1.456 | 0.246 |
| | | | | | React x Day F (1.86, 37.11) = 0.658 | 0.513 |
| | | | | | Treatment x React x Day F (1.86, 37.11)=1.684 | 0.201 |

**Supplemental Table 6: Statistical comparisons: Supplemental Figure 1-con't**

| Figure | Measure | Groups | N-size | Test | F | p value |
|---|---|---|---|---|---|---|
| Suppl T5 | NR vs FR3 PR: Inactive lever presses | NR Veh | 7 | 3-way RM ANOVA | Treatment F (1, 20) = 2.090 | 0.164 |
| | | NR ABC | 6 | | React F (1, 20) = 0.177 | 0.678 |
| | | FR3 Veh | 5 | | Day F (1.95, 38.97) = 0.073 | 0.925 |
| | | FR3 ABC | 6 | | Treatment x React F (1, 20) = 0.492 | 0.491 |
| | | | | | Treatment x Day F (1.95, 38.97) = 1.047 | 0.359 |
| | | | | | React x Day F (1.95, 38.97) = 3.362 | **0.046** |
| | | | | | Treatment x React x Day F (1.95, 38.97)=0.963 | 0.389 |
| Suppl 1Q | Sucrose PR: Active lever presses | Veh | 9 | 2-way RM ANOVA | Treatment F (1, 12) = 1.396 | 0.2602 |
| | | ABC | 5 | | Day F (1.752, 21.02) = 10.87 | **0.0008** |
| | | | | | Treatment x Day F (2, 24) = 0.1143 | 0.8925 |
| Suppl T5 | Sucrose PR: Sucrose pellets | Veh | 9 | 2-way RM ANOVA | Treatment F (1, 12) = 1.409 | 0.2581 |
| | | ABC | 5 | | Day F (1.923, 23.08) = 12.17 | **0.0003** |
| | | | | | Treatment x Day F (2, 24) = 0.3160 | 0.7320 |
| Suppl T5 | Sucrose PR: Inactive lever presses | Veh | 9 | 2-way RM ANOVA | Treatment F (1, 12) = 0.02930 | 0.8669 |
| | | ABC | 5 | | Day F (1.254, 15.05) = 1.054 | 0.3392 |
| | | | | | Treatment x Day F (2, 24) = 1.262 | 0.3012 |
