## Supplemental Table 7 for "Perineuronal nets in the rat medial prefrontal cortex alter hippocampal-prefrontal oscillations and reshape cocaine self-administration memories"

**Supplemental Table 7. Statistical comparisons: Figure 3**

| Figure | Measure | Groups | # animals, (# cells) | Test | F | p value | Šídák's | p value |
|---|---|---|---|---|---|---|---|---|
| 3E | Cell intensity: All PV cells | Veh 1 hr | 4 (746) | 2-way ANOVA | Time F (1, 2420) = 2.986 | 0.0841 | | |
| | | ABC 1 hr | 4 (546) | | Treatment F (1, 2420) = 36.78 | **< 0.0001** | | |
| | | Veh 3 d | 3 (541) | | Time x Treatment F (1, 2420) = 2.884 | 0.0896 | | |
| | | ABC 3 d | 3 (561) | | | | | |
| 3F | PV cell number: All PV cells | Veh 1 hr | 4 | 2-way ANOVA | Time F (1, 10) = 0.02956 | 0.8669 | | |
| | | ABC 1 hr | 4 | | Treatment F (1, 10) = 0.4490 | 0.518 | | |
| | | Veh 3 d | 3 | | Time x Treatment F (1, 10) = 0.8217 | 0.386 | | |
| | | ABC 3 d | 3 | | | | | |
| 3G | Cell intensity: PV/WFA or PV/Stub cells | Veh 1 hr | 4 (296) | 2-way ANOVA | Time F (1, 826) = 0.5724 | 0.4495 | | |
| | | ABC 1 hr | 4 (206) | | Treatment F (1, 826) = 4.171 | **0.0414** | | |
| | | Veh 3 d | 3 (162) | | Time x Treatment F (1, 826) = 4.653 | **0.0313** | 3 days: PV/WFA vs. PV/Stub | **0.0082** |
| | | ABC 3 d | 3 (166) | | | | | |
| 3H | PV cell number: PV/WFA or PV/Stub cells | Veh 1 hr | 4 | 2-way ANOVA | Time F (1, 10) = 0.3839 | 0.5494 | | |
| | | ABC 1 hr | 4 | | Treatment F (1, 10) = 4.392 | 0.0625 | | |
| | | Veh 3 d | 3 | | Time x Treatment F (1, 10) = 0.5477 | 0.4763 | | |
| | | ABC 3 d | 3 | | | | | |
| Suppl 2C | GAD 65/67 puncta | Veh 1 hr | 4 (16) | 2-way ANOVA | Time F (1, 83) = 4.817 | **0.031** | | |
| | | ABC 1 hr | 4 (22) | | Treatment F (1, 83) = 3.118 | 0.0811 | | |
| | | Veh 3 d | 3 (18) | | Time x Treatment F (1, 83) = 0.1751 | 0.6767 | | |
| | | ABC 3 d | 3 (31) | | | | | |
| Suppl 2D | vGLUT1 puncta | Veh 1 hr | 4 (16) | 2-way ANOVA | Time F (1, 83) = 23.22 | **< 0.0001** | | |
| | | ABC 1 hr | 4 (22) | | Treatment F (1, 83) = 0.04075 | 0.8405 | | |
| | | Veh 3 d | 3 (18) | | Time x Treatment F (1, 83) = 0.3049 | 0.5823 | | |
| | | ABC 3 d | 3 (31) | | | | | |
| Suppl 2E | GAD 65/67 to vGLUT1 ratio | Veh 1 hr | 4 (16) | 2-way ANOVA | Time F (1, 78) = 10.52 | **0.0017** | | |
| | | ABC 1 hr | 4 (22) | | Treatment F (1, 78) = 1.390 | 0.2420 | | |
| | | Veh 3 d | 3 (18) | | Time x Treatment F (1, 78) = 1.085 | 0.3007 | | |
| | | ABC 3 d | 3 (31) | | | | | |
