## Supplemental Table 8 for "Perineuronal nets in the rat medial prefrontal cortex alter hippocampal-prefrontal oscillations and reshape cocaine self-administration memories"

**Supplemental Table 8. Statistical comparisons: Figure 4**

| Figure | Measure | Groups | #animals (#cells) | Test | F | p value | Šidák's | p value |
|---|---|---|---|---|---|---|---|---|
| 4C | Excitability | Veh 1 hr | 7 (15) | 3-way RM ANOVA | Current injection F (2.819, 138.1) = 628.7 | **<0.0001** | | |
| | | ABC 1 hr | 4 (12) | | Time F (1, 49) = 2.797 | 0.1008 | | |
| | | Veh 3 d | 10 (17) | | Treatment F (1, 49) = 2.050 | 0.1586 | | |
| | | ABC 3 d | 7 (9) | | Current injection x Time F (6, 294) = 2.565 | **0.0195** | | |
| | | | | | Current injection x Treatment F (6, 294) = 1.797 | 0.0995 | | |
| | | | | | Time x Treatment F (1, 49) = 0.2804 | 0.5988 | | |
| | | | | | Current injection x Time x Treatment F (6, 294) = 0.4164 | 0.8680 | | |
| 4E | 1 hr Firing Variability | Veh 1 hr: 200 pA | 7 (13) | 2-way ANOVA | Current injection F (1, 47) = 0.9415 | 0.3368 | | |
| | | Veh 1 hr: 500 pA | 7 (15) | | Treatment F (1, 47) = 2.284 | 0.1374 | | |
| | | ABC 1 hr: 200 pA | 4 (11) | | Current injection x Treatment F (1, 47) = 0.1734 | 0.6790 | | |
| | | ABC 1 hr: 500 pA | 4 (12) | | | | | |
| 4F | 3 d Firing Variability | Veh 3 d: 200 pA | 10 (17) | 2-way ANOVA | Current injection F (1, 25) = 0.0007894 | 0.9778 | | |
| | | Veh 3 d: 500 pA | 10 (17) | | Treatment F (1, 25) = 0.4675 | 0.5004 | | |
| | | ABC 3 d: 200 pA | 7 (10) | | Current injection x Treatment F (1, 25) = 0.1409 | 0.7106 | | |
| | | ABC 3 d: 500 pA | 7 (10) | | | | | |
| 4G | Capacitance | Veh 1 hr | 7 (16) | 2-way ANOVA | Time F (1, 49) = 0.5931 | 0.4449 | | |
| | | ABC 1 hr | 4 (11) | | Treatment F (1, 49) = 7.415 | **0.0089** | | |
| | | Veh 3 d | 10 (17) | | Time x Treatment F (1, 49) = 0.1575 | 0.6932 | | |
| | | ABC 3 d | 7 (9) | | | | | |
| 4H | Input Resistance | Veh 1 hr | 7 (15) | 2-way ANOVA | Time F (1, 49) = 13.73 | **0.0005** | | |
| | | ABC 1 hr | 4 (12) | | Treatment F (1, 49) = 1.560 | 0.2176 | | |
| | | Veh 3 d | 10 (17) | | Time x Treatment F (1, 49) = 0.7465 | 0.3918 | | |
| | | ABC 3 d | 7 (9) | | | | | |
| 4I | Resting Membrane Potential (RMP) | Veh 1 hr | 7 (15) | 2-way ANOVA | Time F (1, 49) = 2.509 | 0.1196 | | |
| | | ABC 1 hr | 4 (12) | | Treatment F (1, 49) = 0.1801 | 0.6731 | | |
| | | Veh 3 d | 10 (17) | | Time x Treatment F (1, 49) = 3.433 | 0.0699 | | |
| | | ABC 3 d | 7 (9) | | | | | |

### Supplemental Table 8. Statistical comparisons: Figure 4 - con't

| Figure | Measure | Groups | #animals (#cells) | Test | F | p value | Šídák's | p value |
|---|---|---|---|---|---|---|---|---|
| 4J | Action Potential Half-Width | Veh 1 hr | 7 (15) | 2-way ANOVA | Time F (1, 49) = 0.5090 | 0.4790 | | |
| | | ABC 1 hr | 4 (12) | | Treatment F (1, 49) = 0.9509 | 0.3343 | | |
| | | Veh 3 d | 10 (17) | | Time x Treatment F (1, 49) = 0.01987 | 0.8885 | | |
| | | ABC 3 d | 7 (9) | | | | | |
| 4K | Action Potential Threshold | Veh 1 hr | 7 (15) | 2-way ANOVA | Time F (1, 49) = 3.808 | 0.0567 | | |
| | | ABC 1 hr | 4 (12) | | Treatment F (1, 49) = 5.984 | **0.0181** | | |
| | | Veh 3 d | 10 (17) | | Time x Treatment F (1, 49) = 0.4718 | 0.4954 | | |
| | | ABC 3 d | 7 (9) | | | | | |
| 4L | Action Potential Amplitude | Veh 1 hr | 7 (15) | 2-way ANOVA | Time F (1, 49) = 5.170 | **0.0274** | | |
| | | ABC 1 hr | 4 (12) | | Treatment F (1, 49) = 0.2201 | 0.6411 | | |
| | | Veh 3 d | 10 (17) | | Time x Treatment F (1, 49) = 0.7042 | 0.4054 | | |
| | | ABC 3 d | 7 (9) | | | | | |
| 4M | Afterhyper-polarization Amplitude | Veh 1 hr | 7 (15) | 2-way ANOVA | Time F (1, 49) = 1.477 | 0.2300 | | |
| | | ABC 1 hr | 4 (12) | | Treatment F (1, 49) = 2.140 | 0.1499 | | |
| | | Veh 3 d | 10 (17) | | Time x Treatment F (1, 49) = 0.03555 | 0.8512 | | |
| | | ABC 3 d | 7 (9) | | | | | |
| 4N | 1st Interspike Interval | Veh 1 hr | 7 (15) | 2-way ANOVA | Time F (1, 49) = 2.288 | 0.1368 | | |
| | | ABC 1 hr | 4 (12) | | Treatment F (1, 49) = 1.669 | 0.2024 | | |
| | | Veh 3 d | 10 (17) | | Time x Treatment F (1, 49) = 9.689 | **0.0031** | 1 hr Veh vs. ABC | **0.0046** |
| | | ABC 3 d | 7 (9) | | | | ABC: 1hr vs 3d | **0.0093** |
| 4O | Rheobase | Veh 1 hr | 7 (15) | 2-way ANOVA | Time F (1, 49) = 12.99 | **0.0007** | | |
| | | ABC 1 hr | 4 (12) | | Treatment F (1, 49) = 0.08539 | 0.7714 | | |
| | | Veh 3 d | 10 (17) | | Time x Treatment F (1, 49) = 1.270 | 0.2652 | | |
| | | ABC 3 d | 7 (9) | | | | | |
