## Supplemental Table 9 for "Perineuronal nets in the rat medial prefrontal cortex alter hippocampal-prefrontal oscillations and reshape cocaine self-administration memories"

**Supplemental Table 9. Statistical comparisons: Supplemental Figure 3**

| Figure | Measure | Groups | N-size (# trials) | Test | t-test | p value |
|---|---|---|---|---|---|---|
| Suppl 1C | mPFC NR2B bands | FR1 Surface | 6 | Student's t-test | t=0.9087, df=11 | 0.3830 |
| | | VR5 Surface | 7 | | | |
| | | FR1 Intracellular | 6 | | t=0.4248, df=11 | 0.6792 |
| | | VR5 Intracellular | 7 | | | |
| Suppl 1E | Amygdala NR2B bands | Veh 1 hr | 6 | Student's t-test | t=1.249, df=11 | 0.2377 |
| | | ABC 1 hr | 7 | | | |
| | | Veh 3 d | 6 | | t=0.7662, df=11 | 0.4597 |
| | | ABC 3 d | 7 | | | |
