## Supplemental Table 10 for "Perineuronal nets in the rat medial prefrontal cortex alter hippocampal-prefrontal oscillations and reshape cocaine self-administration memories"

### Supplemental Table 10. Statistical comparisons: Figures 5 & 6

| Figure | Measure | Groups (VR session) | N-size (# trials) | Test | t, F or z score | p value | Šídák's | p value |
|---|---|---|---|---|---|---|---|---|
| 5B, C, E | 45-65 Hz Power 200-400 msec | Veh: Last day train | 5 (70) | Student's t-test | Veh: Last day train vs. VR5 (Rewarded) t = 2.001 | **0.0487** | | |
| | | Veh: VR5 | 5 (16) | | | | | |
| 5B, C, F | 35-45 Hz Power 400-450 msec | Veh: Last day train | 5 (70) | Student's t-test | Veh: Last day train vs. VR5 (Rewarded) t = 3.109 | **0.0026** | | |
| | | Veh: VR5 | 5 (16) | | | | | |
| 5B, C, G | 15-30 Hz Power 400-600 msec | Veh: Last day train | 5 (70) | Student's t-test | Veh: Last day train vs. VR5 (Rewarded) t = 2.803 | **0.0063** | | |
| | | Veh: VR5 | 5 (16) | | | | | |
| 5B,C | 2-4 Hz Power 300-700 msec | Veh: Last day train | 5 (70) | Student's t-test | Veh: Last day train vs. VR5 (Rewarded) t = 1.425 | 0.1580 | | |
| | | Veh: VR5 | 5 (16) | | | | | |
| 5C, D, H | 45-65 Hz Power 200-400 msec | Veh, Rewarded | 5 (16) | 2-way ANOVA | Treatment (Veh vs ABC) F (1, 170) = 2.180 | 0.1417 | | |
| | | ABC, Rewarded | 3 (17) | | Lever (Rewarded vs Unrewarded) F (1, 170) = 4.142 | **0.0434** | | |
| | | Veh, Unrewarded | 5 (70) | | Treatment x Lever F (1, 170) = 5.271 | **0.0229** | Veh: Rewarded vs Unrewarded | **0.0058** |
| | | ABC, Unrewarded | 3 (71) | | | | | |
| 5C, D, I | 35-45 Hz Power 400-450 msec | Veh, Rewarded | 5 (16) | 2-way ANOVA | Treatment (Veh vs ABC) F (1, 170) = 16.67 | **< 0.0001** | Rewarded: Veh vs ABC | **< 0.0001** |
| | | ABC, Rewarded | 3 (17) | | Lever (Rewarded vs Unrewarded) F (1, 170) = 0.0778 | 0.7358 | | |
| | | Veh, Unrewarded | 5 (70) | | Treatment x Lever F (1, 170) = 11.37 | **0.0009** | Veh: Rewarded vs Unrewarded | **0.0232** |
| | | ABC, Unrewarded | 3 (71) | | | | ABC: Rewarded vs Unrewarded | 0.0552 |
| 5C, D, J | 15-30 Hz Power 400-600 msec | Veh, Rewarded | 5 (16) | 2-way ANOVA | Treatment (Veh vs ABC) F (1, 170) = 3.819 | 0.0523 | | |
| | | ABC, Rewarded | 3 (17) | | Lever (Rewarded vs Unrewarded) F (1, 170) = 0.7863 | 0.3765 | | |
| | | Veh, Unrewarded | 5 (70) | | Treatment x Lever F (1, 170) = 4.129 | **0.0437** | Rewarded: Veh vs ABC | 0.0556 |
| | | ABC, Unrewarded | 3 (71) | | | | | |
| 5C, D | 2-4 Hz Power 300-700 msec | Veh, Rewarded | 5 (16) | 2-way ANOVA | Treatment (Veh vs ABC) F (1, 170) = 3.960e-005 | 0.9950 | | |
| | | ABC, Rewarded | 3 (17) | | Lever (Rewarded vs Unrewarded) F (1, 170) = 2.136 | 0.1457 | | |
| | | Veh, Unrewarded | 5 (70) | | Treatment x Lever F (1, 170) = 1.902 | 0.1697 | | |
| | | ABC, Unrewarded | 3 (71) | | | | | |

**Supplemental Table 10. Statistical comparisons: Figures 5 & 6 - con't**

| Figure | Measure | Groups (VR session) | N-size (# trials) | Test | t, F or z score | p value | Šídák's | p value |
|---|---|---|---|---|---|---|---|---|
| 6D | Veh Last day train Rewarded | 5-7/70-90 Hz | 4 (12) | Permutation test | Modulation index = 1.92 x 10-4; z = -0.77 | | | |
| 6E | Veh VR5 Rewarded | 5-7/70-90 Hz | 4 (12) | Permutation test | **Modulation index = 6.59 x 10-3; z = 2.12** | | | |
| 6F | ABC VR5 Rewarded | 5-7/70-90 Hz | 3 (17) | Permutation test | **Modulation index = 3.43 x 10-3; z = 1.89** | | | |
| 6H | Granger Prediction Rewarded | Veh Rewarded | 4 (12) | 2-way RM ANOVA | Treatment F (1, 27) = 4.892 | **0.0356** | Veh vs ABC "Press" | **0.0010** |
| | | ABC Rewarded | 3 (17) | | Time F (1, 27) = 6.818 | **0.0146** | | |
| | | | | | Treatment x Time F (1, 27) = 13.20 | **0.0012** | | |
| 6J | Granger Prediction Unrewarded | Veh Unrewarded | 4 (59) | 2-way RM ANOVA | Treatment F (1, 128) = 1.052 | 0.3069 | | |
| | | ABC Unrewarded | 3 (71) | | Time F (1, 128) = 0.7086 | 0.4015 | | |
| | | | | | Treatment x Time (1,128) = 0.0814 | 0.7758 | | |
| 6K, L | Theta lag cross-correlation | Veh Rewarded | 4 (12) | | Peak correlation at 43.5 ± 0.03 ms | | | |
| | | ABC Rewarded | 3 (17) | | Peak correlation at -0.5 ± 0.003 ms | | | |
| 6O | Theta cross-correlation lag | Veh Rewarded | 4 (12) | Kolmogorov-Smirnov | Veh vs ABC, D = 0.6078 | **0.0111** | | |
| | | ABC Rewarded | 3 (17) | | | | | |
| 6P | Theta lag cross-correlation | Veh Rewarded | 4 (12) | 2-way ANOVA | Treatment F (1, 54) = 1.582 | 0.2138 | Veh: Negative vs positive lag | **0.0052** |
| | | ABC Rewarded | 3 (17) | | Lag (positive/negative) F (1, 54) = 5.241 | **0.0260** | ABC: Negative vs positive lag | 0.9754 |
| | | | | | Treatment x Lag F (1, 54) = 6.477 | **0.0138** | | |
