## Supplemental Table 11 for "Perineuronal nets in the rat medial prefrontal cortex alter hippocampal-prefrontal oscillations and reshape cocaine self-administration memories"

**Supplemental Table 11. Statistical comparisons: Supplemental Figure 4**

| Figure | Measure | Groups (VR session) | N-size (# trials) | Test | t, F or z score | p value |
|---|---|---|---|---|---|---|
| Suppl 4C, D | dHIP/mPFC theta/gamma | Veh 8-10/50-70 Hz | 4 (59) | Permutation testing | Modulation index = 4.89 x 10-4; z = 0.24 | |
| | | ABC 8-10/50-70 Hz | 3 (71) | | Modulation index = 1.50 x 10-4; z = -0.86 | |
| Suppl 4E, F | dHIP/mPFC theta cross-correlation lag | Veh | 4 (59) | | Peak correlation at 34.5 ± 0.005 ms | |
| | | ABC | 3 (71) | | Peak correlation at 1.5 ± 0.003 ms | |
| Suppl G | dHIP/mPFC theta cross-correlation lag | Veh | 4 (59) | 2-way ANOVA | Treatment F (1, 256) = 9.736 | **0.0020** |
| | | ABC | 3 (71) | | Lag (positive/negative) F (1, 256) = 1.646 | 0.2006 |
| | | | | | Treatment x Lag F (1, 256) = 2.529 | 0.1130 |
| Suppl H | Median lag at peak correlation | Veh | 4 (59) | Kolmogorov-Smirnov | Veh vs ABC | **0.0028** |
| | | ABC | 3 (71) | | | |
| Suppl 4K, L | Cross-hemispheric mPFC theta | Veh 8-10/70-90 Hz | 5 (70) | Permutation testing | **Modulation index = 2.38 x 10-3; z = 3.18** | |
| | | ABC 8-10/70-90 Hz | 3 (71) | | Modulation index = 1.67 x 10-4; z = -0.71 | |
| Suppl 4M, N | Cross-hemispheric mPFC theta | Veh 5-7/70-90 Hz | 5 (70) | Permutation testing | Modulation index = 1.62 x 10-3; z = 0.41 | |
| | | ABC 5-7/70-90 Hz | 3 (71) | | **Modulation index = 3.04 x 10-3; z = 4.81** | |
| Suppl 4O, P | Cross-hemispheric mPFC theta cross-correlation lag | Veh | 5 (70) | | Peak correlation at -5.5 ± 0.004 ms | |
| | | ABC | 3 (71) | | Peak correlation at -4.5 ± 0.003 ms | |
| Suppl 4Q | Cross-hemispheric mPFC theta cross-correlation lag | Veh | 5 (70) | 2-way ANOVA | Treatment F (1, 232) = 0.4428 | 0.5064 |
| | | ABC | 3 (71) | | Lag (positive/negative) F (1, 232) = 10.57 | **0.0013** |
| | | | | | Treatment x Lag F (1, 232) = 0.02528 | 0.8738 |
| Suppl 4R | Median lag at peak correlation | Veh | 5 (70) | Kolmogorov-Smirnov | Veh vs ABC, D = 0.1645 | 0.4025 |
| | | ABC | 3 (71) | | | |
